# Of bone and ivory: Eastern Atlantic sources of the medieval European walrus ivory trade discovered by ancient mitochondrial and nuclear DNA

**DOI:** 10.64898/2026.08.07.743312

**Authors:** Bastiaan Star, Oliver Kersten, Lydia Hildebrand Furness, Katrien Dierickx, Mohsen Falahati-Anbaran, Anders Söderberg, Natalia Khamaiko, Maja Krzewińska, Anders Götherström, James H. Barrett

## Abstract

Previous molecular archaeological research has made the surprising discovery that almost all studied European walrus finds dating between the 11th and 14th centuries CE seemed to be traded from the Norse colony of Greenland. But why has almost no walrus ivory from the Barents Sea region been detected in medieval Europe, despite its closer proximity and being mentioned in historical accounts? Here, we show that the full spatial extent of medieval Atlantic walrus hunting has been underestimated in previous analyses that targeted modified walrus skulls (rostra) with which pairs of attached tusks were traded, because destructive sampling of ivory artefacts is not always practicable. By analysing workshop offcuts from medieval Sigtuna, Sweden, we directly compare the geographical sources of walrus ivory with rostra, and show that isolated tusks were traded in different ways. We first resolve the genome-wide trans-Atlantic walrus population structure, discovering significant genome-wide nuclear differentiation between western and eastern Atlantic walrus populations. By employing spatially diagnostic nuclear SNPs (*n* = 144,986), we then use low-coverage sequence data to classify historical walrus rostrum and tusk specimens to either western or eastern Atlantic origin, and thus overcome previous limitations when provenancing walruses based on mitochondrial DNA alone. We find that all medieval walrus rostra are assigned exclusively towards the western Atlantic. In contrast, half of the medieval ivory specimens from Sigtuna are assigned to eastern Atlantic sources including Iceland and the Barents Sea region. Moreover, the objects of eastern origin precede those from Norse Greenland in time, suggesting sequential exploitation. Our observations resolve a discrepancy between earlier molecular inference and historical evidence and imply a significantly broader extent and impact of medieval ecological globalisation in the Arctic.

## Introduction

The trade of Arctic walrus ivory in the Middle Ages is among the earliest known examples of ecological globalisation, a process whereby increasingly distant natural resources are exploited on an intercontinental scale, leading to profound socioecological consequences (Barrett et al. 2020). Sequential depletion of walruses in Iceland (Keighley et al. 2019) and western Greenland (Barrett et al. 2020) was driven by demand in Western Europe (Williamson, 2010; Dectot, 2018), Eastern Europe (Barrett et al. 2022; Smirnova 2001; 2005) and perhaps also Byzantium and Asia (Barrett et al. 2022 and references therein). A combination of ancient DNA (aDNA), stable isotope, archaeological and historical evidence together indicates that walruses from increasingly distant sources were hunted, concurrent with a decrease in the size of animals taken (Star et al. 2018; Keighley et al. 2019; Barrett et al. 2020; 2022; Ruiz-Puerta et al. 2024). In addition to impacting walruses, the search for (and sometimes over-exploitation of) this resource has been interpreted as influencing migration (e.g. the settlement of Iceland and Greenland), long-range trade, cultural contact between Norse and Inuit in the High Arctic and potentially the Norse abandonment of Greenland (Frei et al. 2015; Gulløv 2016; Keighley et al. 2019; Arneborg 2021; Barrett 2021; Ruiz-Puerta et al. 2024). Walruses reproduce slowly, gather in large groups, and are therefore especially sensitive to overexploitation (Stewart 2002; McLeod et al. 2014), explaining their rapid extirpation after the Viking Age settlement of Iceland (Keighley et al. 2019), and the serial exploitation of increasingly remote walrus populations in western Greenland (and perhaps, via trade with Inuit, even the eastern Canadian Arctic) by the medieval Norse (Barrett et al. 2020, Ruiz-Puerta et al. 2024). Accurately reconstructing the extent of the historical walrus ivory trade is therefore central to our understanding of early ecological impacts in the Arctic and its place in early globalisation.

Surprisingly, it remains unclear whether walruses of the Barents Sea region (in what is now northern Norway and Russia) were also hunted extensively to supply medieval Eurasian trade. Despite the greater accessibility of the Barents Sea to Europeans, no medieval walrus specimens have previously been attributed to this source based on aDNA, whereas many have been sourced to western Greenland and the eastern Canadian Arctic (Star et al. 2018; Barrett et al. 2020; 2022; Ruiz-Puerta et al. 2024), or in one instance to Iceland (Keighley et al. 2019). A single possible exception from Bergen, Norway (WLR043) has been tentatively attributed to the Barents Sea based on stable isotope evidence, but its aDNA signature is ambiguous (Barrett et al. 2020). The apparent paucity of medieval specimens from the European Arctic is all the more inexplicable given that the earliest known historical evidence for Scandinavian trade of walrus ivory relates to the transport of tusks from the Barents Sea region to Anglo Saxon England, where they were gifted to King Alfred the Great shortly before 900 CE by a northern Norwegian chieftain known as Ohthere or Ottar (Bately 2007; Allport 2025). The potential continued importance of Arctic European sources is further evidenced by the substantial quantities of walrus ivory recovered from medieval (10th to 15th century) Novgorod, a major trading centre with extensive connections across northern Fennoscandia and Russia (Smirnova 2001; 2005).

Here, we ask whether the trade of walrus ivory was more geographically extensive than indicated by previous biomolecular evidence. Did it include the Barents Sea region as well as Iceland, western Greenland and the eastern Canadian Arctic? To answer this question, we test the hypothesis that walrus ivory from Arctic Europe was sometimes traded as isolated tusks rather than as pairs of tusks attached to the front of walrus skulls (rostra). So far, biomolecular research regarding the trade of walrus ivory has focused predominantly on the origin of archaeological walrus rostra (Star et al. 2018, Barrett et al. 2020, Barrett et al. 2022; Ruiz-Puerta et al. 2024). These rostra comprise the front section of the walrus skull, including the bone of the tusk sockets (Figure 1a). Such rostra, sometimes extensively modified and decorated, traveled as a “package” with the ivory tusks they enclosed (Star et al. 2018, Barrett et al. 2020). Upon arrival at processing centres (often towns in Scandinavia, or with Scandinavian connections), the tusks were removed from the bone for local ivory working or onward trade. With an abundant representation in the archaeological record, walrus rostra provide a practical proxy for biomolecular sampling instead of ivory. For instance, one avoids the destructive sampling of precious archaeological ivory carvings by targeting these larger and less carefully modified bone specimens. However, reconstructing ivory trade using rostra hinges on the extent to which they serve as a faithful proxy of all potential sources, an assumption in need of testing by also studying ivory itself. Although it is not always appropriate or practicable to destructively sample ivory artefacts, we here avoid these limitations by studying workshop offcuts of walrus tusk (Figure 1b-e) and minimally invasive samples from artefacts, focusing on a case study from medieval Sigtuna, Sweden. In addition to providing offcuts of tusk ivory, Sigtuna is ideal because rostra have also been found there (allowing direct comparison) (Barrett et al. 2020), it has a time-series of archaeological walrus finds from around 1000 CE until the 13th century CE (Karlsson 2016; Söderberg 2021; Edberg et al. 2022), and it has archaeological evidence for trade connections to both Norway with its North Atlantic network and also (albeit to a lesser extent) Eastern Europe (Tesch 2016; Callmer et al. 2024). Sigtuna thus provides an ideal location to investigate the potential trade of specimens of both Greenlandic/Canadian and European Arctic origin, be the latter traded from Iceland, from northern Fennoscandia through Scandinavian networks, or from what is now the Russian Arctic, via Eastern European networks.

**Figure 1.**
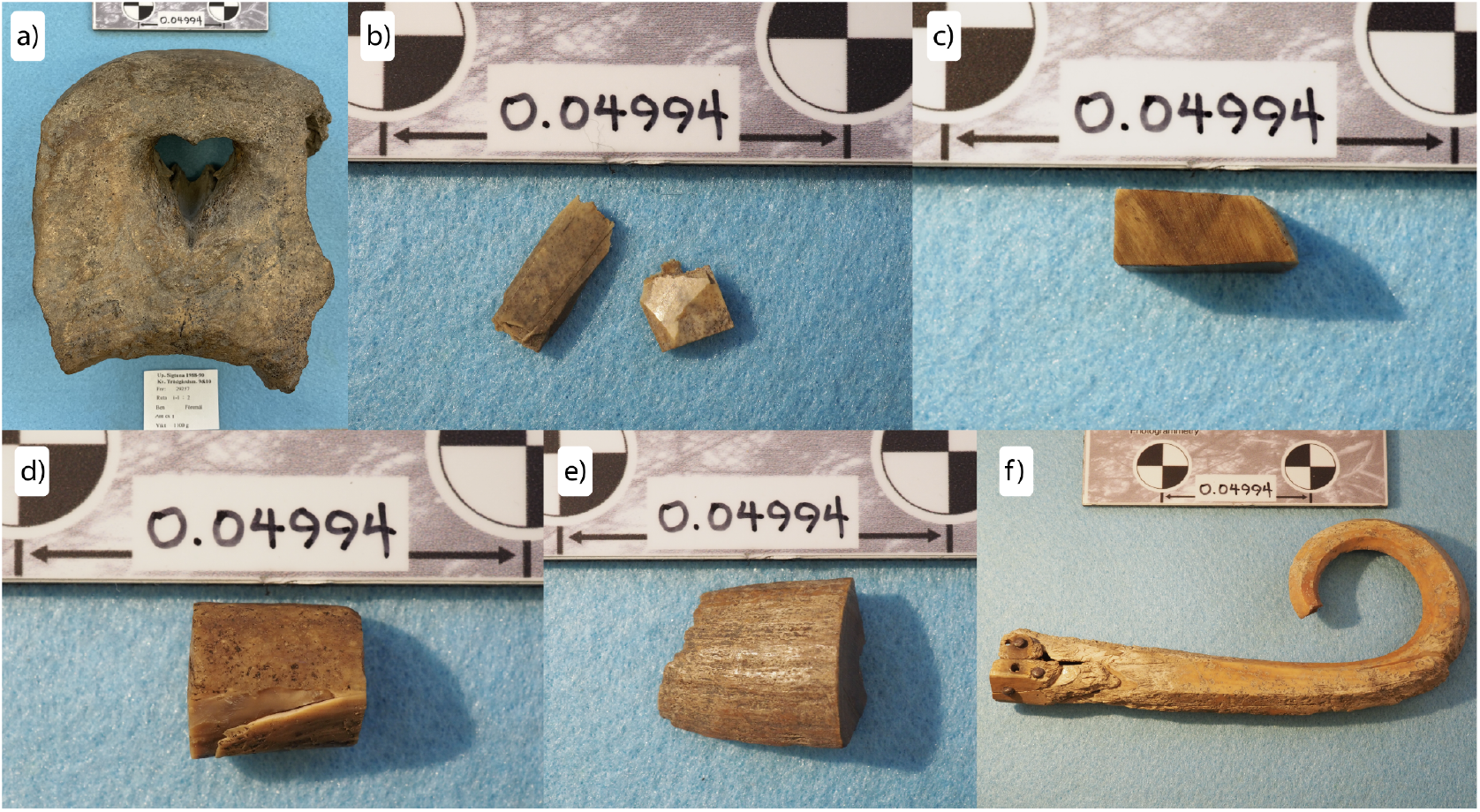
Examples of archaeological bone and ivory specimens of Atlantic walrus from medieval Sigtuna investigated in this study. **a)** A rostrum bone specimen from excavations at kv Trädgårdsmästare (WLR067), **b)** Two pieces of tusk ivory workshop debris (offcuts) from excavations at kv Trädgårdsmästaren 9-10 (WLR082 & WL083), **c)** One tusk ivory offcut from excavations at Professorn 1 (WLR085), **d)** One tusk ivory offcut from excavations at Professorn 1 (WLR088), **e)** One tusk ivory offcut from excavations at Kv. Urmakaren 1 (WLR092), **f)** Head of a crosier from excavation of a probable bishop’s burial at S:ta Gertrud 3 (WLR126). Scale bar ca. 50mm. Full details of each specimen, including museum numbers and dating, are provided in Table S1. (Photos by J.H. Barrett)

Different molecular and morphological approaches have proven effective in assessing the geographical origin of archaeological and historical walrus specimens. First, mitochondrial (mtDNA) analyses enable the assignment of walrus specimens towards specific geographical regions (Star et al 2018, Keighley et al. 2019, Ruiz-Puerta et al. 2024). In particular, it is possible to distinguish a “western clade” present only in western Greenland and Canada, a “north-western” clade present only in north-west Greenland, and an “eastern clade” that is found mainly in the European Arctic, eastern Greenland and previously also Iceland, but also occurs in the western Atlantic (in western Greenland and Arctic Canada) (Star et al. 2018; Ruiz-Puerta et al. 2023). Following the recent publication of additional control data from walruses having known catch locations, it is now also possible to recognise mitogenomic subclades with varying spatial integrity; some of these mtDNA subclades are diagnostic for specific biological sources, whereas others have wider distributions and therefore offer a coarser geographical resolution (Ruiz-Puerta et al. 2024; Dierickx et al. 2026). Second, stable isotope analyses (including δ^13^C, δ^15^N, non-exchangeable δ^2^H and δ^34^S) of walrus rostra can help differentiate specimens coming from western or eastern Atlantic populations (Barrett et al. 2020: 2022). However, isotopic approaches cannot always provide definitive provenance for each specimen, as signatures may be influenced by temporal environmental change (cf. Clark et al. 2022), ecological variation between sexes, and some overlap in baseline data between potential source regions (Dierickx et al. 2026). Third, morphological analysis has identified a distinct manufacturing sequence (*chaîne opératoire*) modifying walrus rostra for trade purposes (Barrett et al. 2020). Because such modified rostra have been recovered directly from excavations in Greenland, these modifications point to a biological origin. Yet the limited number of such archaeological finds introduces important caveats, as their absence elsewhere may reflect gaps in the record rather than a true absence. Together, these approaches are complementary, collectively overcoming the limitations inherent in any single line of evidence, yet are only applicable to rostra as an integrated analytical framework. Collectively, these analyses consistently indicate that medieval rostra found in Europe derived from walrus populations in Greenland and perhaps the eastern Canadian Arctic, traded via the Norse colonies of Greenland (Barrett et al. 2020; 2022; Ruiz-Puerta et al. 2024).

Because the joint transport of rostra holding ivory tusks represents a specific mode of trade, it does not necessarily imply that all walrus ivory reached Europe exclusively via Greenland. If tusks were also traded without being attached to rostra, as is plausible, additional or alternative supply routes cannot be excluded. The Greenlandic signal should therefore be understood as characteristic of the rostrum-based trade, while the full diversity of ivory exchange mechanisms may have been more complex. It has been suggested, for example, that one previously analysed ivory sample from Sigtuna (WLR063, Star et al. 2018) was from a mtDNA subclade that is predominantly confined to Iceland (Keighley et al. 2019). Yet by focusing on the origin of walrus rostra so far, any potential of the trade of isolated tusks from the Barents Sea region has been unaccounted for. Here, we investigate whether the European Arctic has been an underappreciated source for the medieval Atlantic walrus ivory trade by comparing the genetic origin of walrus rostra *vis-a-vis* isolated ivory specimens. As we cannot fully rely on either isotopic or morphological analyses for the ivory itself, we aim to increase the spatial resolution for population assignment of walrus specimens using ancient DNA. We therefore include low-coverage nuclear ancient DNA data, expanding on the previous genetic research by ourselves and others, which has used mtDNA alone (Star et al. 2018; Keighley et al. 2019; Barrett et al. 2020; 2022; Ruiz-Puerta et al. 2024).

Crucially, by including nuclear DNA (nDNA) it becomes possible to obtain an additional line of evidence, resolving any uncertainty whether a specific mtDNA clade derived from the European Arctic (here including eastern Greenland/Iceland in addition to the Barents Sea region) or from western Greenland/Arctic Canada. We describe these alternative geographical origins as the “eastern Atlantic” and “western Atlantic” walrus populations respectively. By further including the most recent mtDNA control data (Ruiz-Puerta et al. 2023; 2024; Dierickx et al. 2026) we then identify a significant number of ivory specimens that are likely to be from Iceland or the Barents Sea region. Finally, we interpret the consequences of our findings for understanding regional resource exploitation and long-distance exchange networks in the medieval North Atlantic.

## Materials and Methods

### Medieval walrus ivory and rostrum samples from Sigtuna

For Sigtuna specimens, we combine ancient DNA data from one previously analysed medieval walrus rostrum (WLR067) and three previously analysed tusk offcuts (WLR063, WLR064 and WLR065) (Star et al. 2018) with results from new analyses of two additional rostra (WLR087, WLR094), nine additional tusk offcuts (WLR082, WLR083, WLR084, WLR085, WLR086, WLR088, WLR090, WLR092, WLR093) and two artefacts of tusk ivory (WLR091, WLR126) (Table S1). The three rostra collectively date between 1160 and 1230 CE based on archaeological context (Karlsson 2016; Table S1); two are represented by recognisable skull bone and one (WLR087) by a piece of cheek tooth. The tusk offcuts are variously dated between 1000 and 1200 CE by archaeological context (Söderberg 2021; Table S1). Of the sampled artefacts, one (WLR091) was a broken fragment of a larger object, perhaps a gaming piece, dated to 1200 to 1230 CE by context (Table S1) and one was a carefully sampled crosier from a probable bishop’s burial, dated between the 11th and early 13th century CE, also based on context (O’Meadhra 2001; Karlsson 2016; Zachrisson and Ljung 2026). Overall, there are 14 specimens of tusk ivory and three from rostra. The find locations of the objects are informative. The five tusk offcuts from excavations at kv Urmakaren (collectively dating between 1000 and 1050 CE) were from part of a royal mint complex (Ros 2009; Söderberg 2021). Moreover, the two rostra and one object of tusk ivory from phase 9 (1200 to 1230 CE) of the kv Trädgårdsmästaren excavations were found near a workshop used for precious metals, glass and rock crystal (Karlsson 2016; Anders Söderberg pers comm.). Combined with the evidence of the crosier noted above, it can thus be inferred that walrus ivory was of high value in Sigtuna throughout the chronology represented by the material.

### Medieval European rostrum samples: comparisons

The 17 specimens from Sigtuna are compared with aDNA data from 30 previously analysed European finds of medieval walrus rostra (Star et al. 2018; Barrett et al. 2020; 2022; Dierickx et al. 2025, Table S1). The rostra range in date from the 11th to 14th/15th centuries CE (Barrett et al. 2020; 2022; Philippsen et al. 2026). Most were excavated from Scandinavian towns or centres with known Scandinavian trade contacts (from Dublin in the west to Kyiv in the east (Barrett et al. 2020; 2022)), although one (WLR073) is without a secure context. As outlined above, the majority of these rostra have been studied using a combination of ancient DNA, stable isotope and manufacturing sequence (*chaîne opératoire*) analysis, with 27 attributed to western Greenland or the eastern Canadian Arctic as a result (Barrett et al. 2020; 2022). Of the remaining three, one (WLR043) was previously tentatively attributed to the Barents Sea region based on stable isotope data, one (WLR104 from Utrecht) underwent aDNA analysis for a different purpose (Dierickx et al. 2025) but is attributed to a source population in the present study and one (WLR040) did not previously provide adequate ancient DNA data (Barrett et al. 2020) but has now been resequenced. Previously analysed rostra with insufficient aDNA preservation are excluded from the present work.

### Control samples from known locations

In addition to the traded European finds of walrus rostra and tusk ivory, a number of natural history and archaeological specimens have been analysed to provide control data from walrus populations in eastern Canada, Greenland, Iceland and the Barents Sea region (Table S1; Table S3). From our own work, these specimens comprise 39 known-origin control samples for mtDNA analysis (Star et al. 2018; Barrett et al. 2020; 2022; Dierickx et al. 2025; 2026; Table S1) and eight known-origin control samples for nDNA analysis (Table S1; Table S2); note that an additional 10 nDNA control samples are drawn from the European archaeological material, as explained below. In addition, we draw on mtDNA data for 89 known-origin control samples of varying date previously published by Keighley et al. (2019) and Ruiz-Puerta et al. (2023; 2024) (Table S3). No nuclear data for these latter specimens is available. Most of the known-origin control specimens are from sites very near to where the walruses would have been hunted. However, a few exceptions were transported within their broad regions of origin. Four specimens are from Igaliku in southwestern Greenland, the location of the medieval bishop’s seat of Gardar (Garðar) to which they must have been transported from further north in western Greenland or the eastern Canadian Arctic (Barrett et al. 2020). One walrus tusk specimen (WLR095, perhaps originally from Svalbard) is from an early modern shipwreck site off the island of Smøla in Central Norway. It is newly published here.

### DNA – Laboratory Work, Sequencing and Mapping

Approximately 300 mg of bone or ivory material was used for ancient DNA (aDNA) analyses for the newly analysed walrus specimens. All pre-PCR procedures were conducted in a dedicated clean room facility at the University of Oslo, Norway, following stringent protocols established for aDNA research (Cooper & Poinar, 2000; Gilbert et al. 2005). Sample surface decontamination was achieved via ultraviolet (UV) irradiation prior to pulverization. Samples were ground into a coarse powder using a sterilized stainless steel mortar and pestle, following established protocols (Gondek et al. 2018). DNA extraction was performed on ∼100 mg of sample powder using a modified pre-digestion protocol (Boessenkool et al. 2017; Lord et al. 2022). Initially, samples were treated with 0.5% sodium hypochlorite (bleach) for 10 minutes and rinsed in nuclease-free water; this step was repeated three times to minimize surface contamination. Subsequently, samples were lysed in a buffer containing 0.5 M EDTA (pH 8.0), 1 M urea, and 10 µg/µl proteinase K, and incubated for 48 hours at ambient temperature. DNA was concentrated using Amicon Ultra-30 kDa centrifugal filter units and further purified with MinElute spin columns (Qiagen). Final elution was carried out in 100 µl of preheated (60°C) elution buffer (EB) as described in Star et al. (2014). DNA yields were quantified using the Qubit fluorometric system to inform downstream processing. Indexed sequencing libraries were prepared following the Santa Cruz dual-indexing protocol (Kapp et al. 2021), employing sample-specific P5 and P7 adapters. Libraries were purified using AMPure XP magnetic beads (Beckman Coulter) and sequenced on the Illumina NovaSeq 6000 platform. In addition to the newly processed material (13 new Sigtuna samples, the Smøla tusk, and one new control sample from Sápmi, northern Fennoscandia), 17 previously published specimens (Table S1) were resequenced (see further below) using previous aDNA extracts or residual sample powder and using the same laboratory protocols outlined above.

Raw sequencing reads from 86 specimens – including newly generated specimens (15 *de novo*, 17 re-sequenced) and 54 from previously published datasets (Star et al. 2018; Barrett et al. 2022; Dierickx et al. 2025; 2026) (Table S2) – were processed using the PALEOMIX pipeline (Schubert et al. 2014). Reads were aligned to the *Odobenus rosmarus* reference genome (Oros_1.0_HiC) (Dudchenko et al. 2017, 2018; Foote et al. 2015), following modifications to the reference assembly. Specifically, (i) only the first 17 Hi-C scaffolds were retained, based on chromosomal boundaries defined by Hi-C contact map data (https://www.dnazoo.org/assemblies/odobenus_rosmarus) and a marked decline in scaffold length beyond this point; (ii) autosomal and sex chromosomal scaffolds were annotated based on synteny with the *Zalophus californianus* reference genome (mZalCal1.pri.v2; GCF_009762305.2); and (iii) the mitochondrial genome (NC_004029.2) was appended to the nuclear reference to facilitate mitochondrial read mapping.

### DNA – Mitogenome Analysis

Following mapping to the nuclear and mitochondrial reference assembly, mitochondrial reads were extracted from the 86 resulting BAM files using samtools (Li et al. 2009). Variant calling was performed with GATK v4.4.0 (McKenna et al. 2010) to identify both single nucleotide polymorphisms (SNPs) and invariant sites. The resulting variant call format (VCF) file was subjected to quality filtering using the expression: QD < 2.0 || MQ < 40 || FS > 60.0 || SOR > 3 || MQRankSum < -12.5 || ReadPosRankSum < -8.0. Additional genotype-level filtering was applied to exclude low-confidence calls using bcftools (Danecek et al. 2021) with the expression -e ‘FMT/DP<3 | FMT/GQ<15’. Filtered variants were converted to FASTA format, with undetermined bases encoded as ‘N’.

Of these 86 specimens, 83 had sufficient MT data and were aligned with 89 mitogenomes previously published by Keighley et al. (2019) and Ruiz-Puerta et al. (2023; 2024) (Table S3), using MAFFT v7.505 (Katoh & Standley, 2013). To minimize alignment ambiguities, the hypervariable control region (starting at position 15,455) was excluded and only specimens with ≥90% genome coverage were retained. The final dataset comprised 172 mitochondrial genomes (Table S1; Table S2; Table S3). Phylogenetic relationships were inferred using IQ-TREE v2.2 (Minh et al. 2020). Model selection was conducted with ModelFinder (Kalyaanamoorthy et al. 2017), and the maximum likelihood tree was reconstructed under the best-fitting evolutionary model identified based on the Akaike Information Criterion (AIC).

### DNA – Nuclear Analysis

#### Genetic sexing of specimens

Genetic sexing of all specimens was performed by comparing the fold-coverage ratio of the Y and X chromosomes. A ratio greater than 0.6 is considered male with a minimum of 10,000 mapped reads following Dierickx et al. (2025).

#### Reference Panel Selection for nuclear population assignment

To further assist the genetic assignment of samples into western and eastern Atlantic populations, nuclear variation was analyzed using the BAMscorer (Ferrari et al. 2022), which assigns low-coverage genomes to predefined clusters based on allele frequency differences at spatially diagnostic SNPs. This approach requires a reference panel to identify the SNPs most strongly associated with genetic divergence between focal populations—in this case, western (western Greenland and the Canadian Arctic) and eastern (e.g. Barents Sea) Atlantic walrus populations. To obtain this reference panel, 18 walrus individuals were selected based on their higher than average endogenous DNA content and previously determined western or eastern Atlantic biological origin. We have selected nine specimens with a western Atlantic origin. Of these, five specimens have a known western Atlantic location based on their western MT clade haplotype (Star et al. 2018; Barrett et al. 2022), whereas four specimens have inferred western location based on stable isotope evidence and pattern of rostrum modification (Barrett et al. 2020; 2022). Eight specimens originate from Svalbard (Star et al. 2018). These specimens are used to establish the nDNA signature for walruses of the eastern mtDNA clade that come from the eastern Atlantic. Finally, we included one ivory trade specimen from Sigtuna with a high endogenous DNA content but unknown source location (Table S2).

#### Genotype Likelihoods for population differentiation

To identify population structure and single nucleotide polymorphisms (SNPs) diagnostic of genetic differentiation between western and eastern Atlantic specimens, genotype likelihoods (GLs) were estimated using the probabilistic framework implemented in ANGSD v0.949 (Korneliussen et al. 2014). Prior to GL estimation, a quality control (QC) assessment was performed to define appropriate thresholds for minimum and maximum read depth, as well as the minimum number of individuals required for genotype likelihood computation. GLs were computed for autosomal sites using the following parameters: *-uniqueOnly 1 -remove_bads 1 -minMapQ 25 -minQ 30 -dosnpstat 1 -C 50 –baq 2 -checkBamHeaders 1 -doHWE 1 -HWE_pval 1e-2 -sb_pval 1e-5 -hetbias_pval 1e-5 -skipTriallelic 1 -snp_pval 1e-6 -minInd 10 -setMaxDepth 52 -rmTrans 1 -doMajorMinor 1 -dobcf 1 -doMaf 1 -doCounts 1 -doGeno 1 -doPost 1 -GL 1 -doGlf 2*, with transitions removed. This filtering resulted in genotype likelihoods at 724,928 variant sites. Population structure was then analyzed using principal component analysis (PCA) as implemented in *PCAngsd* (Meisner & Albrechtsen, 2018) and *NGSadmix* (Skotte et al. 2013). *PCangsd* also outputs SNP loadings along each principal component axis. SNPs with the most extreme loadings are the strongest drivers of the PC. For *NGSadmix*, model convergence was checked with 5 iterations for *k* = (1, 2) and 20 iterations for *k* = (3, 4, 5). The optimal K was determined using the ΔK method (Evanno et al. 2005). The results were visualized using R (R Core Team, 2020).

#### SNP Downsampling and Cutoff Determination for Bamscorer Population Assignment

To statistically determine the minimum number of SNPs required for robust assignment of individuals to eastern or western Atlantic populations using BAMscorer, a downsampling framework was employed in combination with varying SNP weight thresholds derived from PCA axis loadings. For each cutoff (corresponding to the top 5%, 10%, 15%, and 20% of loading values), allele frequency files were generated using the 18 reference samples and the specified SNP positions in ANGSD with the following parameters: *-GL 1 -doMajorMinor 4 -doMaf 8 -doCounts 1 -sites divergent_sites*.*txt*. The resulting .*mafs* output files were converted to BAMscorer-compatible .*frq* format (Supplementary Information).

Power analyses were restricted to those 56 samples with a minimum of 500,000 reads mapped to the nuclear reference genome, ensuring sufficient data quality for statistical evaluation (Table S2). For each of these samples, BAM files were randomly downsampled to a range of sequencing depths: 1,000 to 25,000 reads in 1,000-read increments, and 50,000 to 200,000 reads in 25,000-read increments. Twenty replicate subsamples were generated at each depth level per specimen. To establish baseline assignments, each of the 56 samples was first scored using the Bamscorer program with non-downsampled BAM files and the top 20% quantile of PC1 SNP loadings, resulting in assignment to either western or eastern Atlantic populations. Subsequently, the Bamscorer program was run across all downsampling depths, replicates, and SNP loading cutoffs for each sample. Assignment accuracy was evaluated by assessing the number of consistently classified samples per downsampled read-depth levels. The minimum number of reads required for reliable population assignment was then defined as the bin at which at least 95% of assignments were consistent across all samples and replicates. The most efficient SNP subset, requiring comparatively fewer reads and SNPs to achieve robust classification relative to other cutoffs, was then selected for population assignment of the remaining specimens using BAMscorer.

## Results

### Mitochondrial DNA results

We here analyze 172 Atlantic walrus mitochondrial genomes for which at least ≥90% genome coverage is obtained. We retrieve the expected overall phylogeny which consists of three distinct mitogenome (MT) clades that have been defined as a northwest clade (grey), an eastern clade (orange) and a western clade (blue) based on their geographical distribution (Star et al. 2018, Ruiz-Puerta et al. 2023; 2024, Figure 2a, Figure S1). We find that Sigtuna ivory and rostra specimens cluster within either the eastern or western mitogenomic clades (Figure 2b,c). Specifically, four Sigtuna tusk ivory and two Sigtuna rostra specimens (one of which is represented by a cheek tooth) cluster within the western clade, which is exclusively found in the western Atlantic region. Conversely, ten Sigtuna ivory specimens and one Sigtuna rostrum cluster within the eastern clade, which can be found in either the western or eastern Atlantic region (Figure 2c). Of the ten eastern MT clade ivory specimens, four cluster amongst MT haplotypes that have been retrieved from the Barents Sea area (Figure 2c), whereas three ivory specimens cluster amongst haplotypes from Iceland. The other three eastern MT clade Sigtuna ivory specimens, and the eastern MT clade rostrum, cluster amongst MT haplotypes that are known to occur in the western Atlantic region (Figure 2c).

**Figure 2.**
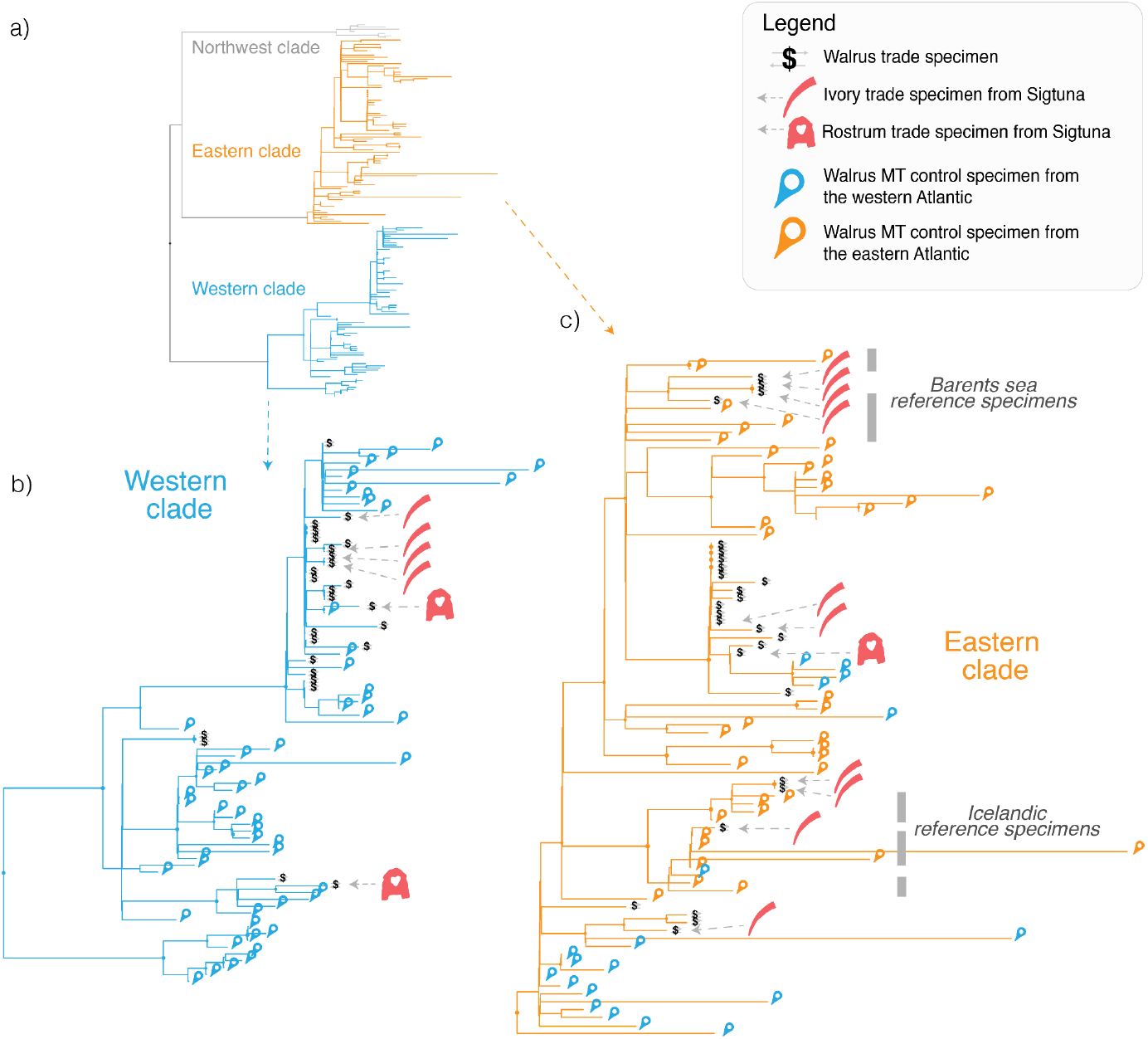
Geographical structure and phylogeny of historical Atlantic walrus mitogenomes. **a)** Three distinct mitogenome (MT) clades segregate within a maximum likelihood tree of 172 Atlantic walrus mitogenomes. These clades have been defined as a northwest clade (grey), an eastern clade (orange) and a western clade (blue) based on the geographical distribution of these clades (Star et al. 2018, Ruiz-Puerta et al. 2023; 2024). **b)** Enlarged detail of the western clade, which is exclusively observed in specimens from the western Atlantic (blue pins), including from western Greenland. Trade specimens with a western clade MT must have come from the western Atlantic (Star et al. 2018), whereas the location of origin of specimens with an eastern clade MT is more ambiguous. **c)** Enlarged detail of the eastern clade, which is mainly observed in specimens from the eastern Atlantic (orange pins), including the Barents Sea region, Iceland and eastern Greenland. Nonetheless, this eastern clade can also be observed in specimens from the western Atlantic (blue pins). Pertinent geographical resolution within the eastern clade is provided by highlighting areas of the phylogeny with reference specimens from the Barents Sea and Iceland (*grey italic text*). Nodes with a circle have greater than 80% support following a Shimodaira-Hasegawa Approximate Likelihood-Ratio Test (SH-aLRT). Sigtuna walrus ivory (red tusk icon) and rostra (red rostra icon) trade specimens ($ icons) occur in the eastern and western clades, as do rostra ($ icons) from elsewhere in Europe. Phylogenetic relationships were inferred using IQ-TREE v2.2 (Minh et al. 2020). Mitogenomic data are from Star et al. (2018), Keighley et al. (2019), Barrett et al. (2022); Ruiz-Puerta et al. (2023; 2024), Dierickx et al. (2025; 2026) and this study (see Table S1; Table S3).

### Nuclear DNA results

We analyze nuclear data of 86 walrus specimens for which we obtained ∼5.7 billion paired reads. We have sufficient data to genetically sex all specimens (Table S2). The dataset is significantly (Binomial exact test: *p* = 0.00032) skewed towards males (*n* = 60) relative to females (*n* = 26). For the 18 specimens that have been included in the walrus reference panel, we analyze ∼4.5 billion paired reads with an endogenous DNA content between 0.08 and 0.65 (Table S2). With this sequencing effort, we obtain a mean ∼2.3 fold nuclear coverage and retrieve 724,928 autosomal nuclear SNPs (excluding transitions) after filtering. These 18 specimens form two distinct groups through visualisation of a PCA analysis (as implemented in *PCAngsd*, Meisner & Albrechtsen, 2018) with the main segregation of specimens following a biomodal clustering pattern along PC1 (Figure 3). Eight specimens from the Barents Sea region cluster together with a single ivory specimen (WLR065) from Sigtuna. Conversely, five rostra specimens with a known western origin (based on their MT genome) cluster together with four other rostra specimens of inferred western origin based on stable isotope evidence and pattern of rostrum modification (Star et al. 2018; Barrett et al. 2020; 2022) (Figure 3). Bimodal population structure was further supported by *NGSadmix*, which identified *K* = 2 as the best model fit in the dataset, with a ΔK value (Evanno et al. 2005) that is multiple orders of magnitude higher than for *K* = 3 or *K* = 4 (Figure S2). Given this distinct bimodal segregation, and considering the associated specimen data, we infer our reference panel to comprise nine specimens (eight from the Barents Sea region and the Sigtuna ivory specimen) that are representative for eastern Atlantic walrus populations and nine specimens (all rostra) that are representative for western Atlantic walrus populations (Figure 3).

**Figure 3.**
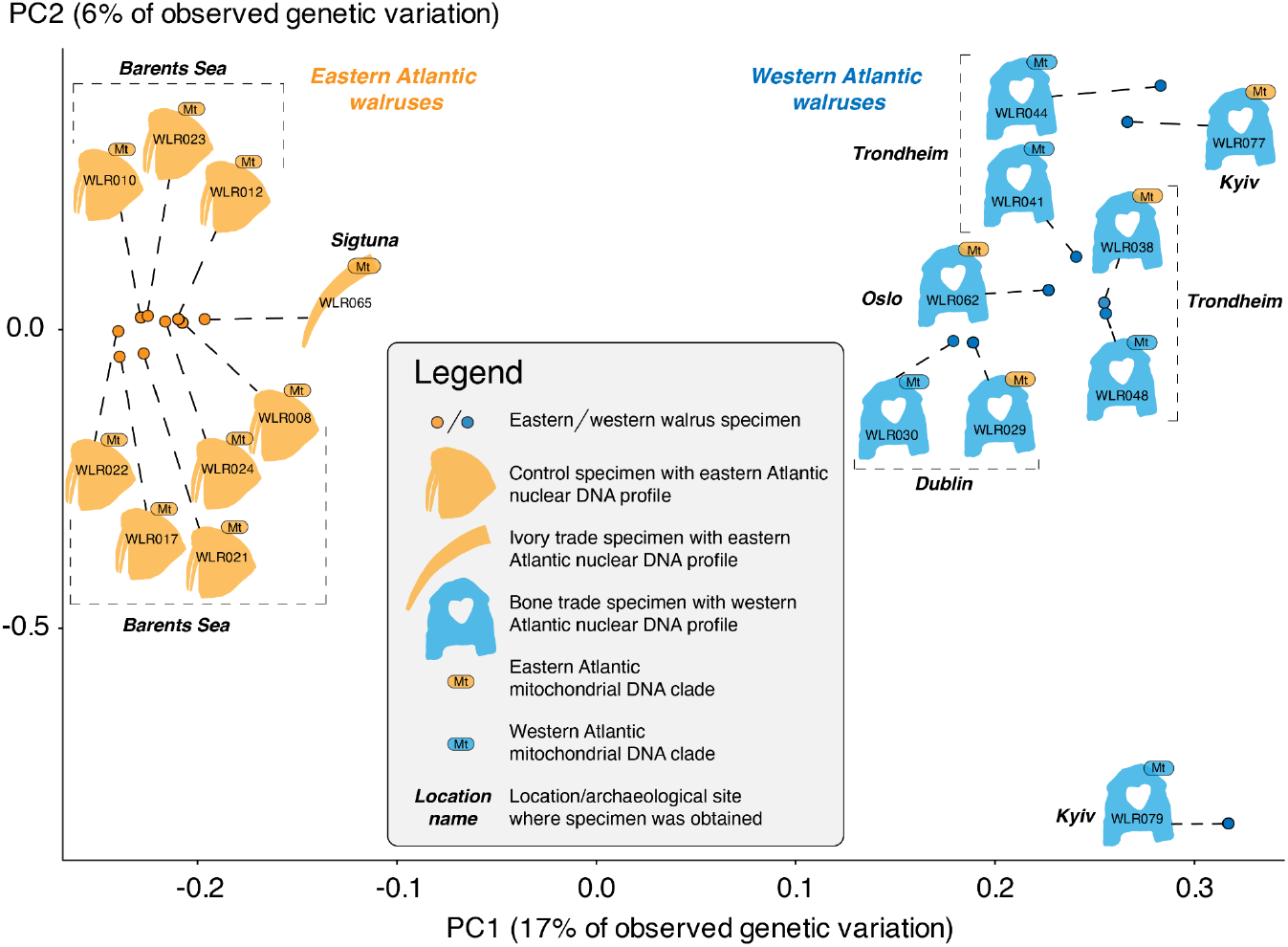
Distinct, bimodal segregation of eastern and western Atlantic walruses revealed through PCA analyses of low-coverage whole genome sequence data. PCA analysis of 18 Atlantic walruses based on 724,928 autosomal nuclear SNPs (excluding transitions) identified two main clusters separated along the first principal component. Eight walruses from the Barents Sea region (orange walrus symbol) cluster with an archaeological ivory trade specimen (orange tusk symbol) from Sigtuna. These nine walruses all have an eastern Atlantic mitochondrial (MT) clade (orange MT oval). Five walrus rostra trade specimens (blue rostrum symbol) have a MT clade that is exclusively confined to the western Atlantic cluster (blue MT oval) and therefore must have originated from the western Atlantic (Star et al. 2018; Barrett et al. 2022). These five rostra specimens cluster together with four other rostra specimens having an eastern-like MT clade (orange MT oval) that has been introgressed into the western Atlantic region (Star et al 2018). We conclude that these four specimens also originate from the western Atlantic, based on co-segregation here and the fact that they were previously inferred to have a western origin based on stable isotope evidence and method of rostrum modification (Barrett et al. 2020; 2022). Diagnostic SNPs (*n* = 144,986) yield a weighted *Fst* divergence of at least 0.12 between western and eastern walruses. Individual specimen identification numbers are plotted inside each symbol.

We then analyse subsets of spatially diagnostic SNPs from this reference panel that drive the bimodal population segregation, to assess our ability to use extremely low-coverage sequencing data to genetically assign archaeological walrus specimens to either western or eastern Atlantic populations. By selecting either the top 5%, 10%, 15%, or 20% of SNP loading values along PC1, we obtain respectively 36247, 72493, 108740 or 144986 SNPs for this subset. These SNPs represent the strongest drivers of population structure between our 18 reference specimens, yielding a weighted Fst divergence of at least 0.12 between the reference specimens.

Power analyses using 56 walrus individuals for which we have obtained at least 500,000 mapped nuclear reads show that the most efficient SNP subset is at a 20% cut-off. This cut-off requires comparatively fewer reads and SNPs to achieve robust classification relative to other cut-offs values (Figure S3). At this 20% cut-off, robust classification (>95% accuracy) is achieved with a minimum of 50,000 mapped reads for eastern Atlantic specimens (Figure 4a), and a minimum 140,000 reads for western Atlantic specimens (Figure 4b). Requiring a minimum number of 140,000 reads for all our specimens, we can confidently assign 80 low coverage walrus specimens to either the western or eastern Atlantic walrus populations (Table S2). Two specimens have between 50,000 and 140,000 reads and have an eastern population classification. These specimens with fewer reads are more ambiguous based on nuclear genetic data only, yet all are control samples of known eastern Atlantic origin (e.g. from the Barents Sea region) so the genetic classification is consistent with the specimen metadata in these cases for which origin is known.

**Figure 4.**
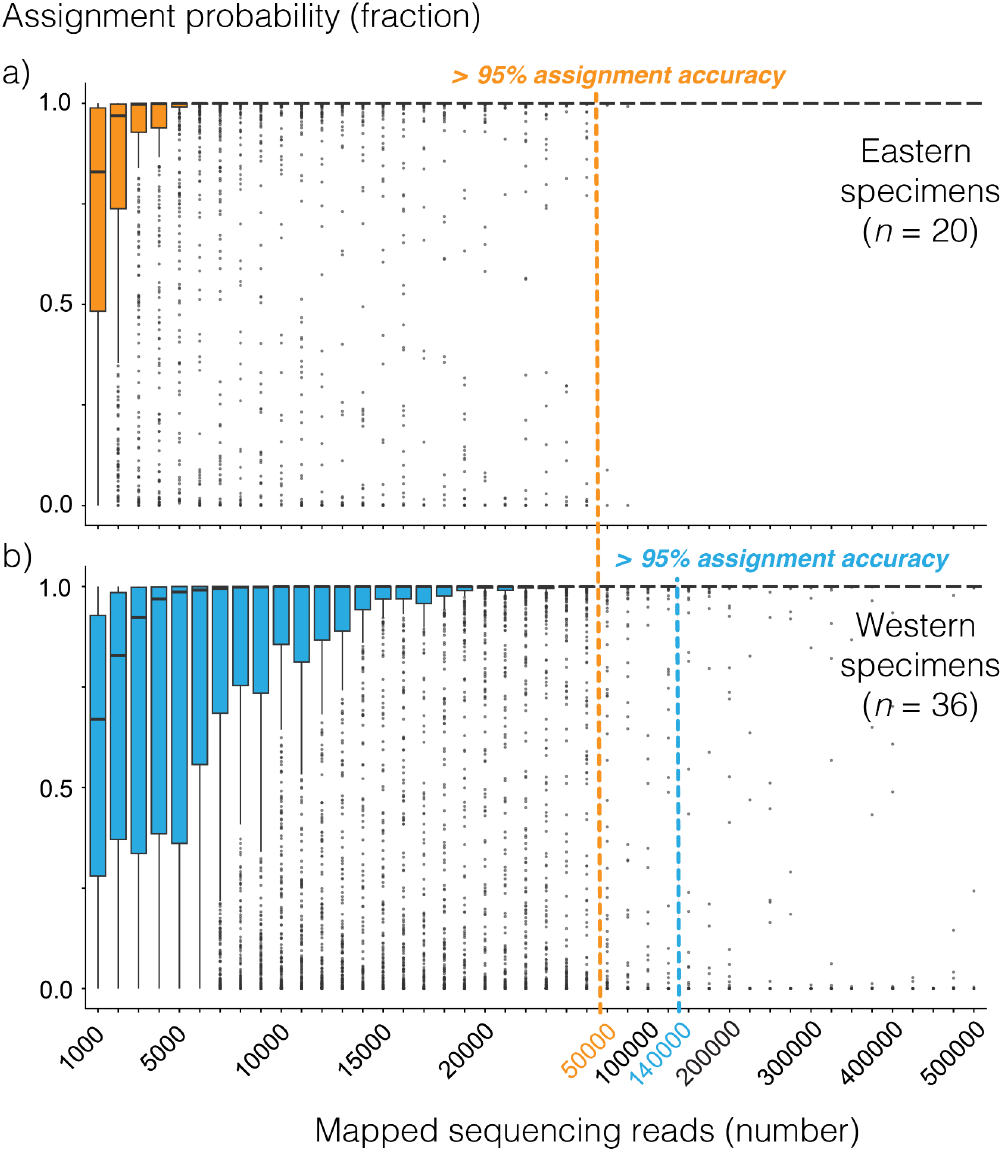
The limits of genetically assigning Atlantic walrus specimens to western or eastern Atlantic populations using extremely low-coverage sequencing data. The accuracy and consistency of population assignment using low-coverage sequencing data is investigated by downsampling reads from Atlantic walrus specimens for which at least 500,0000 sequencing reads have been aligned to the nuclear reference genome. At each read-depth level, random downsampling (without replacement) is iterated (*n* = 20) per specimen. Each iteration is scored using BAMscorer with the assignment probabilities belonging to the eastern or western population plotted as a boxplot. Population assignment is based on the 20% most genetically divergent autosomal SNPs (*n* = 144,986, weighted *Fst* differentiation 0.12) that separate the reference dataset of 18 walrus individuals (see Figure 3). At higher read-depth level (> 500,000 reads) walrus specimens are bimodally assigned to either the eastern or western population. **a)** *Assignment accuracy of 20 eastern walrus specimens*. At 50,000 mapped reads (orange dotted line), more than 95% of iterations have an assignment probability of 1 towards the eastern Atlantic. **b)** *Assignment accuracy of 36 western walrus specimens*. At 140,000 mapped reads (blue dotted line), more than 95% of iterations have an assignment probability of 1 toward the western Atlantic.

### Combined mito-nuclear population assignment

We have sufficient sequencing data for a combined nuclear population assignment of 80 walrus specimens, including the reference specimens used to generate the reference panel (Table S2, Figure 5). All relevant control specimens from northern Canada and Greenland (*n* = 14) are consistently assigned to the western Atlantic population by nuclear classification (Figure 5). Two of the Greenland specimens have the eastern MT clade, which is known to have spread towards western Greenland (Star et al. 2018). Similarly, all control specimens from Svalbard and Norway (*n* = 20) are consistently assigned the eastern Atlantic population by nuclear classification (Figure 5). All of these specimens have an eastern MT clade (if not missing due to lack of data), which is the exclusive MT clade observed in the eastern Atlantic (Star et al. 2018). All archaeological walrus rostra from western Greenland and Europe having sufficient reads (*n* = 34) are exclusively assigned to the western Atlantic population by nuclear classification (Figure 5). These rostra specimens have either a western or eastern MT clade. In contrast, the tusk ivory specimens from Sigtuna (*n* = 14) are assigned to both western and eastern Atlantic populations by nuclear classification. Of these Sigtuna tusk ivory specimens, seven are assigned to the western Atlantic population and they have both western and eastern MT clades. The remaining seven specimens are assigned to the eastern Atlantic population and all have eastern MT clades. The fraction of eastern versus western Atlantic specimens (7/7) is significantly higher in Sigtuna ivory compared to all walrus rostra (0/34, two-tailed Fisher’s Exact test *p* < 0.0001). The Sigtuna eastern population ivory specimens are exclusively males whereas the Sigtuna western population ivory and rostra specimens comprise both males and females, although this pattern is not statistically significant with the present sample sizes (Table S4).

**Figure 5.**
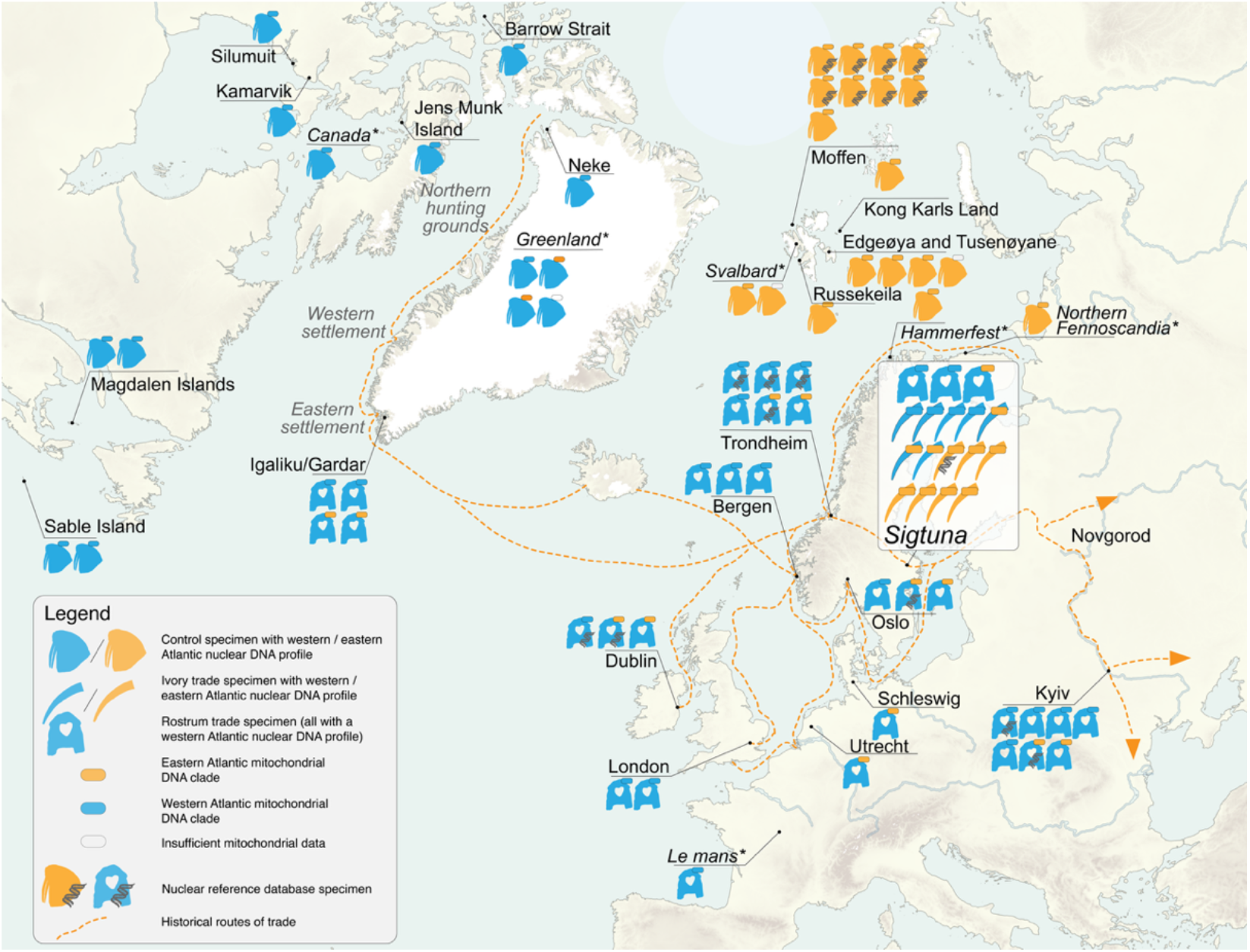
An eastern Atlantic origin of ivory from Viking Age Sigtuna shown by mito-nuclear population assignment of 80 walrus specimens. Well-preserved Atlantic walrus reference specimens with higher endogenous DNA (DNA symbol, *n* = 18) have been used to provide genome-wide divergence data that allows the population assignment of specimens with lower coverage nuclear sequencing data (*n* = 62). Walrus specimens (rostrum or tusk) are genetically assigned as belonging to western (blue) or eastern (orange) Atlantic populations. Specimens are also assigned to an exclusively western (blue oval) or geographically more wide-spread eastern (orange oval) mitogenomic clade. All rostrum trade specimens (blue rostrum symbol, *n* = 34) are assigned to the western Atlantic region using nuclear population assignment. Tusk ivory trade specimens from Sigtuna (blue or orange tusk symbol) are assigned equally to the western Atlantic (*n* = 7) or eastern Atlantic (*n* = 7) walrus population. Nuclear population assignment of control (non-trade) specimens with known locations (blue or orange walrus symbol) is fully consistent with an overall east-west genetic population segregation. For some specimens (*) associated geographic information is an approximate representation, rather than a precise locality. Mitogenomic data are from Star et al. (2018), Keighley et al. (2019), Barrett et al. (2022); Ruiz-Puerta et al. (2023; 2024), Dierickx et al. (2025; 2026) and this study (see Table S1; Table S3; nuclear data are from Star et al. (2018), Barrett et al. (2022); Dierickx et al. (2025; 2026) and this study (see Table S1; Table S2).

## Discussion & Conclusion

Here, we set out to investigate if the trade of walrus ivory tusks was more geographically extensive than indicated by previous biomolecular evidence, which was based on rostrum bone specimens that were likely exclusively sourced from or via Greenland. We start by improving our capacity to source the origin of Atlantic walrus ivory fragments by generating the first genome-wide nuclear data (nDNA) of historic specimens coming from the entire Atlantic region. Significant genetic structure across this range has been observed earlier based on mitogenomic data (Star et al. 2018; Keighley et al. 2019; Ruiz-Puerta et al. 2023; 2024). Specifically, the majority of mitogenomic haplotypes in Atlantic walrus fall within two genetic lineages, one, the “western clade” with a distribution limited to the western Atlantic, and another, the more wide-spread “eastern clade” that is present in the European Arctic, eastern Greenland, previously Iceland and also the western Atlantic (Star et al. 2018; Keighley et al. 2019; Ruiz-Puerta et al. 2023; 2024). The origin of these lineages has been dated to approximately 20,000 years ago (Star et al. 2018; Ruiz-Puerta et al. 2023) which fits a scenario of separation within glacial refugia on either side of the Atlantic Ocean during the last glacial maximum (LGM), followed by secondary contact with directional gene flow from eastern Atlantic walruses into the western Atlantic (Star et al. 2018). Here, we find that significant trans-Atlantic population divergence is also present within the nuclear DNA of Atlantic walruses. This observation has implications for our understanding of Atlantic walrus population structure, migration and our ability to source archaeological walrus specimens.

Because mtDNA is maternally inherited, the previously observed mtDNA structure reflects female dispersal only (Star et al. 2018; Keighley et al. 2019). Thus, as females have more limited home ranges than males (Dietz et al. 2014), male-mediated gene flow could have homogenized allele frequencies at nuclear loci, following expansion northwards and westwards during the early Holocene or more recently. Our results show that such male-mediated gene flow has not been so extensive as to erase all genetic differentiation. Instead, the observed nuclear genetic structure implies that both sexes are philopatric to a degree that trans-Atlantic gene flow has remained sufficiently rare to preserve significant differentiation between walruses from the western Atlantic and eastern Atlantic. These genetic results agree with inference from tagging studies showing that – despite high mobility – walrus males nonetheless exhibit remarkable site fidelity over multiple years (Born et al. 2005; Lydersen & Kovacs, 2014; Mikkelsen et al. 2024). Overall, this consistent genetic divergence between western and eastern Atlantic walrus populations fits a scenario of refugial isolation during the LGM, with subsequent independent demographic histories leading to mitonuclear differentiation on either side of the Atlantic Ocean.

The sourcing capacity of this nuclear east-west DNA divergence, combined with substructure within the mtDNA phylogeny, yields particular insights when applied to the medieval ivory trade. Thirty-four of the 37 medieval walrus rostra considered herein (three from Sigtuna, 30 from elsewhere in Europe and four from Gardar/Igaliku in Greenland) have sufficient DNA preservation for attribution to a source population based on combined nDNA and/or mtDNA analysis. All derive from western walrus populations present in western Greenland and the eastern Canadian Arctic. They must thus have reached Europe via the Norse colony of Greenland, be they hunted in Greenland or in some cases perhaps the Canadian Arctic (Barrett et al. 2020; Ruiz-Puerta et al. 2024). This observation includes 13 rostra of the eastern mtDNA clade, that were of less certain origin without complementary nDNA data. On present evidence, walrus tusks traded to Europe as pairs attached to modified rostra thus represented a specifically Greenlandic product in the Middle Ages. There remains only one known possible exception; WLR043 (a rostrum from Bergen) is consistent with the Barents Sea based on previous stable isotope analysis (Barrett et al. 2020), but its origin is ambiguous based on the present study due to insufficient nDNA reads.

Of the 17 walrus specimens from Sigtuna studied herein, three were rostra (one represented by a cheek tooth that was originally part of a rostrum, given that traded mandibles, which also contain cheek teeth, have not been observed in medieval Europe (Barrett 2021)). As mentioned above, all three were classified as deriving from the western population based on nDNA. The observation that most or all walrus rostra from medieval Europe, including those from Sigtuna, were ultimately derived from the Norse colonies of Greenland corroborates previous interpretations (Barrett et al. 2020: 2022; Ruiz-Puerta et al. 2024). Conversely, the 14 objects of walrus tusk ivory from Sigtuna (two artefacts and 12 workshop offcuts) were evenly divided between specimens derived from the western and eastern Atlantic walrus populations. Based on the mitochondrial maximum likelihood tree, of the seven objects from the eastern population, three are most likely to derive from Iceland and four from the Barents Sea region.

This new evidence clarifies that ivory tusks traded without associated rostra were harvested from both Iceland and the Barents Sea region. The discrepancy between aDNA and historical evidence especially considering the Ohthere account of a northern Norwegian chieftain acquiring walrus tusks from Sámi hunters (Bately 2007; Allport 2025) – is thus resolved, and the broad extent of medieval walrus hunting is better documented. Overall, walrus ivory for medieval European sculpture and craftworking was not only sourced from western Greenland (and probably Arctic Canada), but also from the Barents Sea region and Iceland.

The present results also have broader implications for interpreting the eastern circulation of walrus ivory beyond Scandinavia. The archaeological distribution of walrus finds (ivory and/or rostra), encompassing both Eastern and Southeastern Europe, is attested by finds from Novgorod (Smirnova 2001; 2005), Kyiv (Sahaidak et al. 2008; Sahaidak et al. 2015; Khamaiko 2018; Barrett et al. 2022), Sarkel in the Volga–Don region (Artamonov 1956: Fig. 25) and Dorostolon in the Balkans (Atanasov 1987: Pls. IX:8; X:8), many of which are broadly dated to the 12th century. That the walrus rostra recovered from Kyiv derive from western Atlantic populations is consistent with a shift from eastern to western resource procurement. At the same time, the Kyivan Chronicle records an exchange of princely gifts under the year 1160 CE, in which walrus tusks (described as ‘fish tooth’) are listed together with Arctic fox and white wolf pelts (PSRL 1998: fols. 180–180v). This gift assemblage provides valuable documentary evidence for the circulation of prestigious commodities of potentially northern European origin. Archaeological and historical evidence further indicates that, from the late 11th century onwards, Rus’ of Eastern Europe progressively incorporated the northern peoples of the European Arctic into a sphere of political influence, tribute collection and long-distance exchange, while the trade in northern furs is documented both archaeologically and in the Novgorod birch-bark letters (Makarov 1997; Lapshin 2019). Against this broader historical background, it is an important question for future research whether some walrus ivory circulating in Eastern Europe derived from the Barents Sea region, especially prior to the 12th century.

The ecological consequences of this trade for walruses in the European Arctic remain unknown, as the present case study is based on a relatively small number of ivory fragments from a single settlement. Nonetheless, a tantalizing chronological pattern emerges. It is notable that six of the seven Sigtuna objects of eastern origin predate 1100 CE (with the seventh dating between 1130 and 1160), whereas nine of the ten western walrus specimens (six of tusk ivory and three rostra) all postdate 1100 CE (the latest being from the first half of the 13th century). This chronological change in origin is highly significant (two-tailed Fisher’s Exact test, *p* = 0.0037). During the 1100s, Greenlandic walrus ivory may thus have replaced tusks from the two eastern sources (there being no difference in date between the probable Icelandic and Barents Sea specimens). Such replacement could have resulted partly from new trade networks, for example associated with the establishment of a bishopric in Greenland in the 12th century and subsequent ecclesiastical participation in the ivory trade (Star et al. 2018). However, the medieval extirpation of the Icelandic walrus is immediately pertinent (Keighley et al. 2019), and it should be asked whether the abundance of walruses around the southern shores of the Barents Sea was also impacted, as later occurred in Svalbard after the discovery of that archipelago in 1596 (Kovacs et al. 2014). Pending further research, sequential depletion is a potential explanation of the apparent western shift in the source of ivory used in Sigtuna, as has been observed for the northward progression of walrus hunting in Norse Greenland later in the Middle Ages, during the 13th and 14th centuries (Barrett et al. 2020). This chronological shift therefore hints at broader ecological consequences of medieval ivory exploitation in the European Arctic than previously recognized.

How did the walrus finds reach medieval Sigtuna, a town in the Swedish interior far removed from all of the sources identified by aDNA? The town was well situated to receive long-distance trade goods by land and sea, from north, west and east. Prior to isostatic rebound, trade goods originally from northern Norway, Iceland and Greenland could have arrived by sea via what is now Lake Mälaren (Tesch 2016). Alternatively, the trade may have followed a long-standing east-west land route through Jämtland to Trondheim Fjord in Norway (Hennius 2021; Baug et al. 2024); there, the town of Trondheim, at the crossroads of travel northward to Arctic Norway and westward to Iceland and Greenland, was a focus of 11th-century and later trade in walrus ivory and rostra (Barrett 2021; 2025). For the material post-dating the 11^th^ century, Bergen could also have served as an important transshipment centre (Barrett et al. 2020). A predominantly western facing trade, by land or sea, might best explain the contemporary presence of walrus ivory from Iceland and the Barents Sea in the 11th to 12th centuries, and of predominantly Greenlandic material thereafter. Sigtuna could alternatively have received walrus tusks from Arctic Russia via intermediaries in Eastern Europe, although the town’s connections in this direction were more limited than observed for its predecessor Birka (Callmer et al. 2024). It is also possible that walrus ivory from the Barents Sea could have reached Sigtuna via northern Sweden. Here silver hoard evidence may reflect the wealth of intermediaries trading with Sámi hunters, principally for furs (Androshchuk 2026). Regardless of the specific route or routes, combined mtDNA and nDNA analysis of walrus ivory and rostra from Sigtuna greatly expands our understanding of the spatial extent of medieval walrus hunting, resolves a past inconsistency between biomolecular and historical evidence, improves our understanding of walrus population history, and opens a new methodological window for further historical ecological study of the ivory trade.

## Acknowledgements

Samples and/or associated metadata for previously unpublished results were kindly provided by: Sigtuna Museum, The University Museum of Bergen (Gitte Hansen, Anne Karin Hufthammer, Hanneke Johanna Maria Meijer), Natural History Museum, London (Richard Sabin, Roberto Portela Miguez, Phaedra Kokkini), NTNU University Museum (Axel Christophersen, Torkel Johansen, Birgit Maixner, Staale Normann, Jon Anders Risvaag, Fredrik Skoglund, Birgitte Skar), Museum of Cultural History, University of Oslo (Marianne Vedeler), Institute of Archaeology of the National Academy of Sciences of Ukraine, Kyiv (Mykhailo Kublii), and the National Museum of Ireland, Dublin (Maeve Sikora). Bente Philippsen kindly advised regarding radiocarbon dating. Ole Risbøl kindly transported and cleared Norwegian Customs with many of the Sigtuna specimens. Torun Zachrisson and Mats Roslund kindly assisted with pertinent sources regarding Sigtuna. We thank Danielle Buss, Erin Kunish and Thomas Royle for sampling several of the previously published control samples and for collegial discussions regarding walrus biology and/or archaeological walrus finds. We further acknowledge the Norwegian Sequencing Centre (University of Oslo; https://www.sequencing.uio.no) for the sequencing of genomic libraries. Sequence data computations were performed on resources provided by Sigma2— the National Infrastructure for High-Performance Computing and Data Storage in Norway under project NN9244K and NN11026K.

## Ethics

Permission to sample the previously unpublished archaeological and natural history material was kindly provided by the museums responsible for the relevant specimens (please see Table S1 for a comprehensive list). The newly analysed samples from Sigtuna were transferred internationally to Norway under CITES export permit numbers 4.10.18-15058/2022 and 4.10.18-05121/2024. Because Norway is not a member of the European Union, CITES import permits are not required for walrus, an Appendix III/list C-species; permission to import was thus confirmed by direct correspondence with the Norwegian Environment Agency. The newly analysed sample from the United Kingdom was transferred internationally under institutional CITES registrations GB 001 and NO 007. Where relevant, previously reported samples were shipped across borders using the CITES export and import permits or institutional CITES registrations noted in the articles where first published.

## Data Availability

Nuclear and mitochondrial reads used in this study have been uploaded to the European Nucleotide Archive (ENA) and can be accessed under project PRJEB120774. Mitochondrial control data from Canada, Greenland and Iceland have also previously been published and released by Keighley et al. (2019) and Ruiz-Puerta et al. (2023; 2024).

## Author contributions

Conceptualization: B.S., J.H.B. Resources: A.S., N.K., M.K., A.G. Methodology: B.S., O.K., L.H.F., K.D., M.K., A.G., J.H.B. Investigation: B.S., O.K., J.H.B. Formal analysis: B.S., O.K., J.H.B. Data curation: B.S., O.K, M.F.-A., J.H.B. Visualization: B.S. Writing – original draft: B.S., J.H.B. Writing review & editing: B.S., O.K., L.H.F., K.D., M.F.-A., A.S., N.K., M.K., A.G., J.H.B. Supervision: B.S., J.H.B. Funding acquisition: B.S., J.H.B. All authors gave final approval for publication and agree to be held accountable for the work.

