## Supplementary_information for "Of bone and ivory: Eastern Atlantic sources of the medieval European walrus ivory trade discovered by ancient mitochondrial and nuclear DNA"

**Script to convert ANGSD mafs output files to BAMscorer-compatible.frq format.**

```
#!/bin/bash
# =============================================================================
# Script: Convert ANGSD.mafs.gz output to BAMscorer-compatible .frq format
# Input:  Per-chromosome ANGSD minor allele frequency files (.mafs.gz)
# Output: Population allele frequency files (.frq) for BAMscorer
# =============================================================================
# -----------------------------------------------------------------------------
# STEP 1: Concatenate per-chromosome MAF files into a single autosomal file Chr 1 is used with zcat directly (includes
# header); chrs 2-15 have their headers stripped with tail -n +2 before appending.
# -----------------------------------------------------------------------------
for pop in AA BB; do
    echo ${pop}
    # Decompress chr 1 (retains the header line)
    zcat Walrus_${pop}_chr_1.mafs.gz > Walrus_${pop}_chr_Autosomes.mafs
    # Append chromosomes 2-15, skipping their header lines
    for i in {2..15}; do
        zcat Walrus_${pop}_chr_${i}.mafs.gz | tail -n +2 >> Walrus_${pop}_chr_Autosomes.mafs
    done
    # Count total SNP sites (excluding header); expected ~144,986
    cat Walrus_${pop}_chr_Autosomes.mafs | tail -n +2 | wc -l
done
# Expected output: 144,986 sites per population
# -----------------------------------------------------------------------------
# STEP 2: Clean up MAF values that slightly exceed 1.0 due to floating-point rounding in ANGSD. Any frequency > 1.0 is capped  # at exactly 1.000000. Also strips the header line for downstream awk processing.
# -----------------------------------------------------------------------------
for pop in AA BB; do
    echo ${pop}
    # Strip header line
    tail -n +2 Walrus_${pop}_chr_Autosomes.mafs > Walrus_${pop}_chr_Autosomes_edit.mafs
    # Cap MAF values exceeding 1.0 (column 6) to exactly 1.000000
    awk -F '\t' '{$6 = ($6 > 1.000000 ? "1.000000" : $6)}1' OFS='\t' \
        Walrus_${pop}_chr_Autosomes_edit.mafs > Walrus_${pop}_chr_Autosomes_edit2.mafs
done
# -----------------------------------------------------------------------------
# STEP 3: Fix missing minor allele bases at sites where MAF == 0 When ANGSD estimates a MAF of 0, the minor allele base (col  # 4) is arbitrary/undefined. Here we replace it with the minor allele reported for the same site in the other population's # file, ensuring consistency.
# -----------------------------------------------------------------------------
# For AA: borrow minor allele base from BB where AA MAF == 0
# FNR==NR: read BB file first, storing minor allele (col 4) keyed by chrom+pos
# Then for AA file: if MAF (col 6) == 0, replace its minor allele with BB's
awk -F '\t' 'FNR==NR{a[$1,$2]=$4; next} {if($6==0.000000){$4=a[$1,$2]}}1' OFS='\t' \
    Walrus_BB_chr_Autosomes_edit2.mafs \
    Walrus_AA_chr_Autosomes_edit2.mafs > Walrus_AA_chr_Autosomes_edit3.mafs

# For BB: borrow minor allele base from AA where BB MAF == 0
awk -F '\t' 'FNR==NR{a[$1,$2]=$4; next} {if($6==0.000000){$4=a[$1,$2]}}1' OFS='\t' \
    Walrus_AA_chr_Autosomes_edit2.mafs \
    Walrus_BB_chr_Autosomes_edit2.mafs > Walrus_BB_chr_Autosomes_edit3.mafs
# -----------------------------------------------------------------------------
# STEP 4: Reformat MAF data into PLINK-style.frq format for BAMscorer Output columns: CHROM, POS, N_ALLELES, N_CHR, # AJOR_ALLELE:FREQ, MINOR_ALLELE:FREQ  N_CHR is calculated as N_individuals * 2 (diploid). Major allele frequency = 1 - minor # allele frequency
# -----------------------------------------------------------------------------
for pop in AA BB; do
    echo ${pop}
    cp Walrus_${pop}_chr_Autosomes_edit3.mafs tmp
    # Write the .frq header line (matches PLINK .frq format expected by BAMscorer)
    echo -e "CHROM\tPOS\tN_ALLELES\tN_CHR\t{ALLELE:FREQ}" > ${pop}.frq
    # Reformat each site:
    #   col 1 = chromosome, col 2 = position
    #   col 3 = major allele, col 4 = minor allele
    #   col 6 = minor allele frequency, col 7 = number of individuals
    awk '{
        Chrom=$1; Pos=$2
        N_Alleles=2
        N_Ind=$7; N_Chr=N_Ind*2          # diploid: multiply individuals by 2
        MinAll=$4; MinFreq=$6
        MajAll=$3; MajFreq=1-MinFreq     # major freq is complement of minor freq
        printf("%s\t%s\t%s\t%s\t%s:%.6f\t%s:%.6f\n",
            Chrom, Pos, N_Alleles, N_Chr,
            MajAll, MajFreq, MinAll, MinFreq) >> "'${pop}'.frq"
    }' tmp
    rm tmp  # Clean up temporary file
done
# -----------------------------------------------------------------------------
# STEP 5: Copy final .frq files to output files with descriptive BAMscorer names
# -----------------------------------------------------------------------------
OUT=Walrus_BamscorerRef
cp AA.frq ${OUT}_AA_SNPs.frq
```

cp BB.frq ${OUT}_BB_SNPs.frq

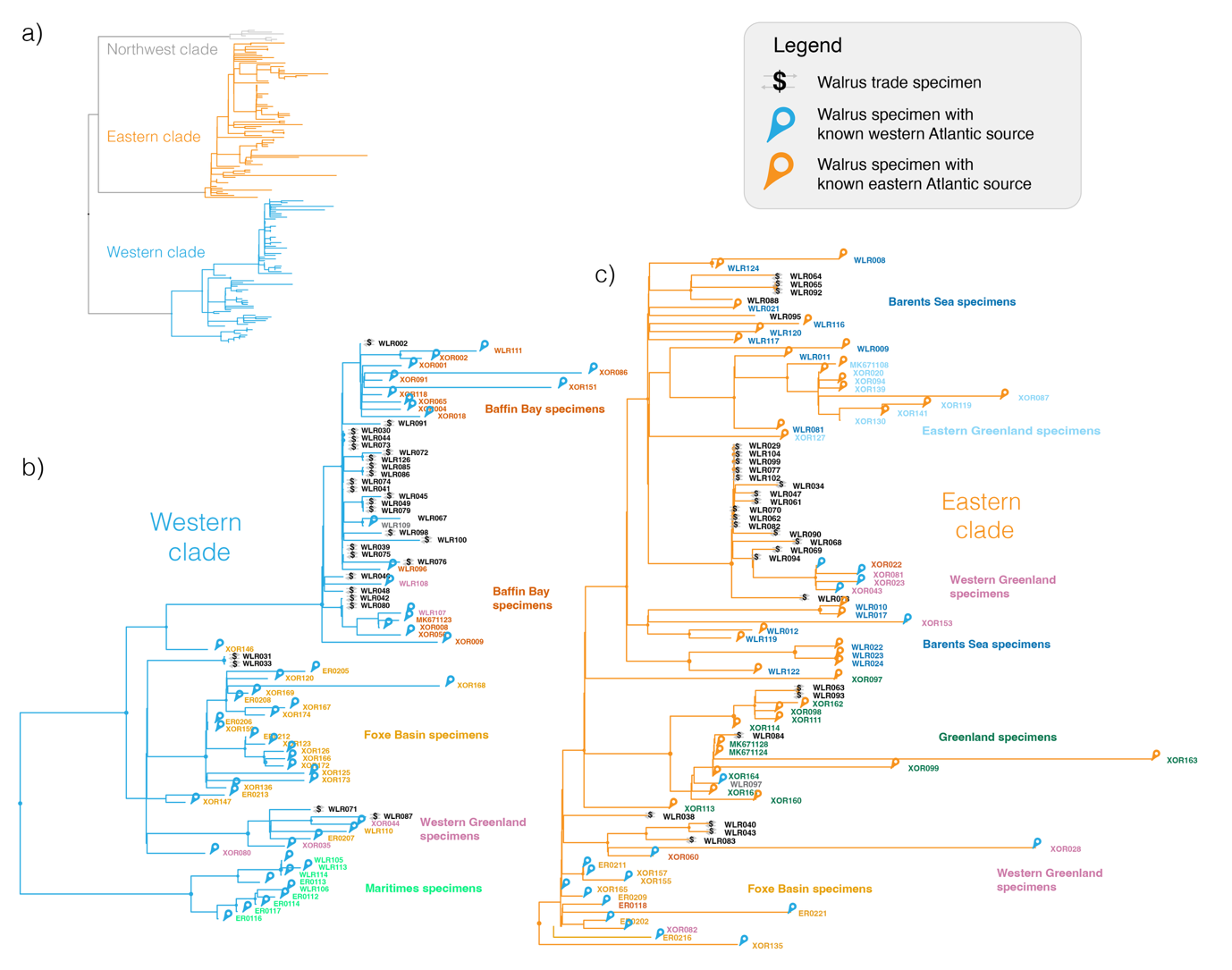
**Supplementary Figure 1.** **Geographical structure and phylogeny of historical Atlantic walrus mitogenomes. a)** Three distinct mitogenome (MT) clades segregate within a maximum likelihood tree of 173 Atlantic walrus mitogenomes. These clades have been defined as a northwest clade (grey), an eastern clade (orange) and a western clade (blue) based on the geographical distribution of these clades (Star et al. 2018, Ruiz-Puerta et al. 2024). **b)**  Enlarged detail of the western clade, which is exclusively observed in specimens from the western Atlantic (blue pins), including from western Greenland. Trade specimens with a western clade MT must have come from the western Atlantic (Star et al. 2018), whereas the location of specimens with an eastern clade MT is more ambiguous. **c)** Enlarged detail of the eastern clade, which is mainly observed in specimens from the eastern Atlantic (orange pins), including the Barents Sea region, Iceland and eastern Greenland. Nonetheless, this eastern clade can also be observed in specimens from the western Atlantic (blue pins). Nodes with a circle have greater than 80% support following a Shimodaira-Hasegawa Approximate Likelihood-Ratio Test (SH-aLRT). The specimen identification number for control specimens colour coded according to their sample origin following Dierickx et al. 2026. Mitogenomic data are from Star et al. (2018), Keighley et al. (2019), Barrett et al. (2022); Ruiz-Puerta et al. (2023; 2024), Dierickx et al. (2025; 2026) and this study (see Table S1; Table S3).

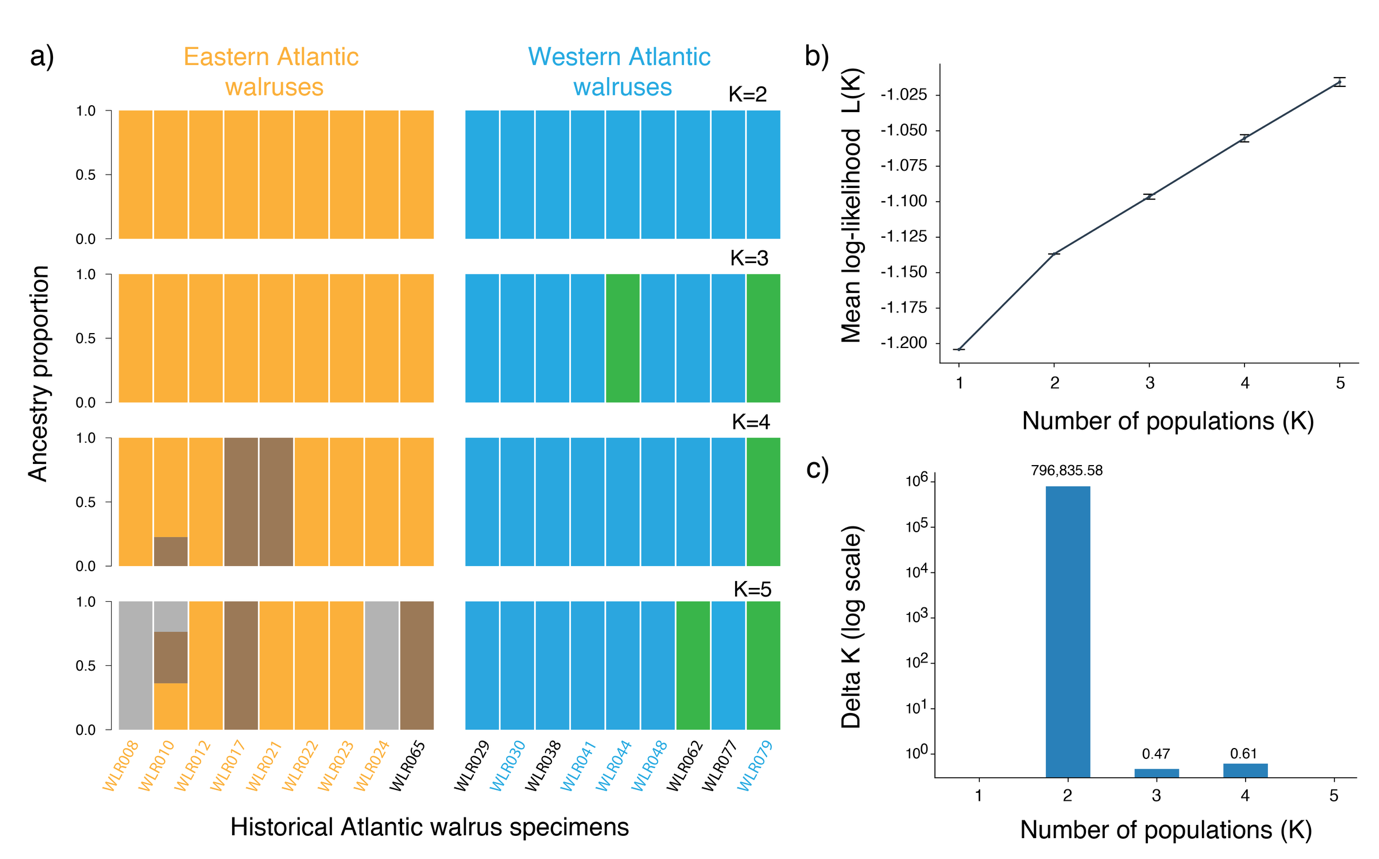

**Supplementary Figure 2. Distinct, bimodal segregation of eastern and western Atlantic walruses revealed through admixture analyses of low-coverage whole genome sequence data. a)** Admixture proportions of 18 historical Atlantic walruses as quantified by *NGSadmix* based on 724,928 autosomal nuclear SNPs (excluding transitions). Coloured bars indicate *NGSadmix* cluster for each individual at each K value (K=2–K=5). Clusters are not matched across K values. At K = 2 we observe two main clusters with eight walruses from the Barents Sea region (WLR008 to WLR024, orange label) clustering with an archaeological ivory trade specimen (WLR065) from Sigtuna (black label). Conversely, five walrus rostra trade specimens (WLR030, WLR041, WLR044, WLR048 and WLR079, blue label) have a MT clade that is exclusively confined to the western Atlantic cluster and therefore must have originated from the western Atlantic (Star et al. 2018; Barrett et al. 2022). Four other specimens (WLR029, WLR038, WLR062, and WLR077, black label) have an eastern-like MT clade that has been introgressed into the western Atlantic region (Star et al. 2018) and segregate with these five western specimens. We conclude that these four specimens also originate from the western Atlantic, based on co-segregation here and the fact that they were previously inferred to have a western origin based on stable isotope evidence and pattern of rostrum modification (Barrett et al. 2020; 2022). See also Figure 3 and text of the main manuscript. **b)** The mean log-likelihood as a function of the number of ancestral populations (K) across replicate *NGSadmix* runs for each value of K (K = 1: *n* = 5; K = 2: *n* = 5; K = 3–5: *n* = 20 each). Error bars represent ±1 standard deviation across replicates and indicate run-to-run convergence at each K; error bars are visually negligible at K = 1–2 but increase with K, reflecting greater variability among independent runs. **c)** Evanno's ΔK statistic across K = 2–4. ΔK (Evanno et al. 2005) was calculated from the mean and standard deviation of log-likelihoods, using the second-order rate of change in log-likelihood standardized by its standard deviation. The pronounced peak at K = 2 (ΔK ≈ 7.97 × 10⁵) shows this is the best model fit compared to K = 3 and K = 4 (ΔK = 0.47 and 0.61, respectively).

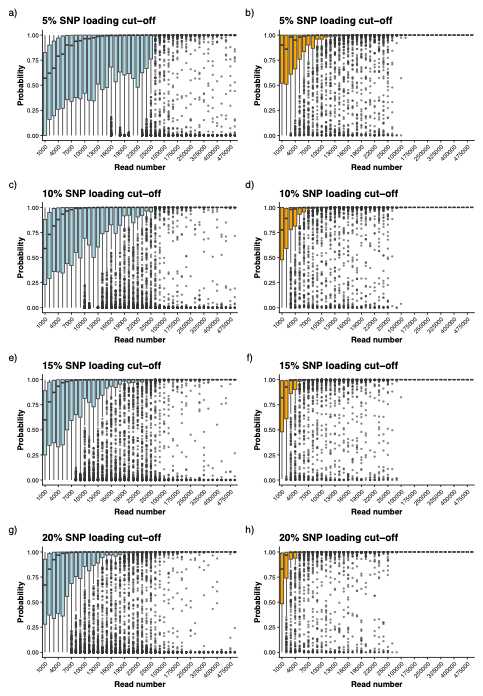

**Supplementary Figure 3. Assessing genetic assignment of Atlantic walrus specimens to western or eastern Atlantic populations using extremely low-coverage sequencing data.** The accuracy and consistency of population assignment using low-coverage sequencing data is investigated by downsampling reads from 56 Atlantic walrus specimens for which at least 500,0000 sequencing reads have been aligned to the nuclear reference genome. At each read-depth level, random downsampling (without replacement) is iterated (*n* = 20) per specimen. Each iteration is scored using BAMscorer (Ferrari et al. 2022) with the assignment probabilities belonging to the western (blue, *n* = 36) or eastern (orange, *n* = 20) population plotted as a boxplot. Population assignment is based on 5% (panel a & b), 10% (panel c & d) 15% (panel e & f) or 20% (panel g & h) most genetically divergent autosomal SNPs. At higher read-depth level (> 500,000 reads) walrus specimens are bimodally assigned to either the eastern or western population.

**Dating**

Where available, Table S1 provides calendrical dates for the specimens employed for combined mitochondrial and nuclear DNA analysis. Most of the traded medieval archaeological tusk ivory and rostrum samples are dated by archaeological context or (in a few instances) stylistic attributes of carved decorations/runic inscriptions. The relevant details are from museum metadata that were supplied when sampling and/or can be found in the references cited in Table S1 and the main text. A few of the medieval samples, and many of the natural history and archaeological control samples, also have published radiocarbon dates (Dierickx et al. 2025; 2026; Philippsen et al. 2026). These have been individually recalibrated for the present work using OxCal Version 4.4 (Bronk Ramsey 2001; Bronk Ramsey 2009), the Marine20 calibration curve (Heaton et al. 2020) and ΔR values from Pieńkowski et al. (2022) for specimens from (or inferred originally to have been from) the Barents Sea region, Philippsen et al. 2026 for specimens from (or inferred originally to have been from) Greenland or the Canadian Arctic, and pertinent molluscan samples from McNeely et al. (2006) as reported in http://calib.org/marine/ (Reimer and Reimer, 2001) for specimens from the Magdalen Islands and Sable Island. Two new radiocarbon dates, for samples WLR081 and WLR095, are published here for the first time and are calibrated as noted above. For several natural history specimens, the historically recorded kill year is known from museum metadata. Moreover, in some instances without a known kill year, the precision of calibrating the radiocarbon dates for natural history specimens was improved by incorporating the historical date of first known museum registration as a terminus post quem (TPQ) using the following OxCal code (this example is for aDNA specimen WLR117):

Sequence()

{

Curve("Marine20","marine20.14c");

Delta_R(-61,37);

Boundary();

R_Date("Tra-23506",575,12);

Before(C_Date("registered",1929));

Boundary();

};
