## Supplementary_tables for "Of bone and ivory: Eastern Atlantic sources of the medieval European walrus ivory trade discovered by ancient mitochondrial and nuclear DNA"

Table S1. Sample details for walrus specimens included in the combined mitochondrial and nuclear DNA analysis.

| aDNA Unique Identifier | Identifiers from 4OCEANS & Northern Journeys Projects | Radiocarbon Lab Number (TRa-) | Resequenced | Latitude | Longitude | Museum | Traded Artefact | Rostrum, Tusk Ivory or Other | Find Location | Country | Museum Number | Data First Published | Archaeological Age (CE) | Historical Age (CE) | Date First Registered if Historical (CE) | Uncalibrated Radiocarbon Date (BP) | Uncalibrated Radiocarbon Date Error | Calibrated Radiocarbon Date BCE/CE Median | Calibrated Radiocarbon Date BCE/CE From (95.4%) | Calibrated Radiocarbon Date BCE/CE To (95.4%) | MT Clade | Available Nuclear Data | Number of Mapped Reads (n) | Sufficient Reads for Western Nuclear Assignment | Sufficient Reads for Eastern Nuclear Assignment | Nuclear Clade (Bamscorer) | Comments | Combined Classification |
|---|---|---|---|---|---|---|---|---|---|---|---|---|---|---|---|---|---|---|---|---|---|---|---|---|---|---|---|---|
| WLR002 | R20 | | No | 60.398 | 5.325 | Bergen University Museum | Yes | Rostrum | Bergen | Norway | JS540_BRM0/81304/b | Barrett et al. 2020 & this study | 1170 to 1198 | NA | NA | | | | | | West | Yes | 152320 | Yes | Yes | West | | West |
| WLR008 | JS1855_B1950 | | Yes | 80.029 | 14.500 | Bergen University Museum | No | Other | Moffen, Svalbard | Norway | JS1855_B1950 | Star et al. 2018 & this study | 17th to 19th century | NA | NA | | | | | | East | Yes | 138420421 | Yes | Yes | East | | East |
| WLR009 | JS366_OT_06 | | No | 78.079 | 13.737 | Bergen University Museum | No | Other | Russekeila, Svalbard | Norway | JS366_OT_06 | Star et al. 2018 | 18th to 19th century | NA | NA | | | | | | East | Yes | 1513126 | Yes | Yes | East | | East |
| WLR010 | JS1855_B1947 | | No | 80.029 | 14.500 | Bergen University Museum | No | Other | Moffen, Svalbard | Norway | JS1855_B1947 | Star et al. 2018 | 17th to 19th century | NA | NA | | | | | | East | Yes | 6531818 | Yes | Yes | East | | East |
| WLR011 | JS1855_B2036 | | No | 80.029 | 14.500 | Bergen University Museum | No | Other | Moffen, Svalbard | Norway | JS1855_B2036 | Star et al. 2018 | 17th to 19th century | NA | NA | | | | | | East | Yes | 1466898 | Yes | Yes | East | | East |
| WLR012 | FOC5089 | 24929 | Yes | 80.029 | 14.500 | Bergen University Museum | No | Other | Moffen, Svalbard | Norway | JS1855_B1952 | Star et al. 2018 & this study | 17th to 19th century | NA | NA | 593 | 16 | 1837 | 1699 | 1953 | East | Yes | 35780426 | Yes | Yes | East | | East |
| WLR017 | JS1855_B2034 | | Yes | 80.029 | 14.500 | Bergen University Museum | No | Other | Moffen, Svalbard | Norway | JS1855_B2034 | Star et al. 2018 & this study | 17th to 19th century | NA | NA | | | | | | East | Yes | 142933996 | Yes | Yes | East | | East |
| WLR021 | FOC3079 | 22913 | Yes | 80.029 | 14.500 | Bergen University Museum | No | Other | Moffen, Svalbard | Norway | JS1855_B1877 | Star et al. 2018 & this study | 17th to 19th century | NA | NA | 699 | 14 | 1737 | 1559 | 1909 | East | Yes | 150582649 | Yes | Yes | East | | East |
| WLR022 | FOC5132 | | Yes | 80.029 | 14.500 | Bergen University Museum | No | Other | Moffen, Svalbard | Norway | JS1855_B1907 | Star et al. 2018 & this study | 17th to 19th century | NA | NA | | | | | | East | Yes | 93360443 | Yes | Yes | East | | East |
| WLR023 | JS1855_B2005 | | No | 80.029 | 14.500 | Bergen University Museum | No | Other | Moffen, Svalbard | Norway | JS1855_B2005 | Star et al. 2018 | 17th to 19th century | NA | NA | | | | | | East | Yes | 4338170 | Yes | Yes | East | | East |
| WLR024 | FOC5124 | | Yes | 80.029 | 14.500 | Bergen University Museum | No | Other | Moffen, Svalbard | Norway | JS1855_B1973 | Star et al. 2018 & this study | 17th to 19th century | NA | NA | | | | | | East | Yes | 139197896 | Yes | Yes | East | | East |
| WLR029 | R31 | | No | 53.344 | -6.277 | National Museum of Ireland | Yes | Rostrum | Dublin | Ireland | E173:4330 | Star et al. 2018 | NA | NA | NA | | | | | | East | Yes | 3669169 | Yes | Yes | West | | West |
| WLR030 | R30 | | No | 53.344 | -6.277 | National Museum of Ireland | Yes | Rostrum | Dublin | Ireland | E173:323 | Star et al. 2018 | NA | NA | NA | | | | | | West | Yes | 3320527 | Yes | Yes | West | | West |
| WLR031 | R32 | | No | 51.516 | -0.092 | MOLA | Yes | Rostrum | London | UK | 21273 | Star et al. 2018 | late 12th to early 13th century | NA | NA | | | | | | West | Yes | 1920148 | Yes | Yes | West | | West |
| WLR033 | R33 | | No | 51.516 | -0.092 | MOLA | Yes | Rostrum | London | UK | 21274 | Star et al. 2018 | late 12th to early 13th century | NA | NA | | | | | | West | Yes | 210538 | Yes | Yes | West | | West |
| WLR034 | R29 | | No | 53.344 | -6.274 | National Museum of Ireland | Yes | Rostrum | Dublin | Ireland | E172:4311 | Star et al. 2018 | 11th century | NA | NA | | | | | | East | Yes | 598889 | Yes | Yes | West | | West |
| WLR038 | FOC5298/R6 | | Yes | 63.431 | 10.400 | NTNU University Museum | Yes | Rostrum | Trondheim | Norway | N10102 | Star et al. 2018 & this study | 1111 or shortly before | NA | NA | | | | | | East | Yes | 27988033 | Yes | Yes | West | | West |
| WLR039 | R10 | | No | 60.398 | 5.325 | Bergen University Museum | Yes | Rostrum | Bergen | Norway | BRM 0/46092/1 | Star et al. 2018 | 1120 to 1170 | NA | NA | | | | | | West | Yes | 87152 | No | Yes | NA | MT west, in MT tree | West |
| WLR040 | R17 | | Yes | 60.396 | 5.327 | Bergen University Museum | Yes | Rostrum | Bergen | Norway | BRM 76/19742/1 | Barrett et al. 2020 & this study | 1250 to 1300 | NA | NA | | | | | | East | Yes | 102003 | No | Yes | NA | MT east, in MT tree | NA |
| WLR041 | FOC4811/R3 | | Yes | 63.431 | 10.401 | NTNU University Museum | Yes | Rostrum | Trondheim | Norway | N32091 | Star et al. 2018 & this study | 1225 to 1275 | NA | NA | | | | | | West | Yes | 61352584 | Yes | Yes | West | | West |
| WLR042 | R12 | | No | 60.399 | 5.323 | Bergen University Museum | Yes | Rostrum | Bergen | Norway | BRM 4/4176/1 | Star et al. 2018 | late 12th century to 1225 | NA | NA | | | | | | West | Yes | 141584 | Yes | Yes | West | | West |
| WLR043 | R8 | | No | 60.398 | 5.325 | Bergen University Museum | Yes | Rostrum | Bergen | Norway | BMR 0/2709/1 | Star et al. 2018 | 1248 to 1332 | NA | NA | | | | | | East | Yes | 80994 | No | Yes | NA | MT east, in MT tree | NA |
| WLR044 | FOC4813/R2 | | Yes | 63.431 | 10.401 | NTNU University Museum | Yes | Rostrum | Trondheim | Norway | N29494 | Star et al. 2018 & this study | 1150 to 1175 | NA | NA | | | | | | West | Yes | 94591655 | Yes | Yes | West | | West |
| WLR045 | R11 | | No | 60.398 | 5.325 | Bergen University Museum | Yes | Rostrum | Bergen | Norway | BRM 0/87411/1 | Star et al. 2018 | 1248-1332 | NA | NA | | | | | | West | Yes | 1430046 | Yes | Yes | West | | West |
| WLR046 | FOC4812/R1 | 25177 | No | 63.426 | 10.396 | NTNU University Museum | Yes | Rostrum | Trondheim | Norway | N167587 | Star et al. 2018; Philippsen et al. 2026 | 1500 to 1532 but residual in context | NA | NA | 1229 | 13 | 1340 | 1236 | 1442 | West | Yes | 283791 | Yes | Yes | West | | West |
| WLR047 | FOC4810/R4 | 25175 | No | 63.431 | 10.401 | NTNU University Museum | Yes | Rostrum | Trondheim | Norway | N37603 | Star et al. 2018; Philippsen et al. 2026 | post 1600 but residual in context | NA | NA | 1389 | 21 | 1194 | 1055 | 1302 | East | Yes | 3420742 | Yes | Yes | West | | West |
| WLR048 | FOC5297/R5 | | Yes | 63.430 | 10.401 | NTNU University Museum | Yes | Rostrum | Trondheim | Norway | N203145 | Star et al. 2018 & this study | 12th century | NA | NA | | | | | | West | Yes | 18113210 | Yes | Yes | West | | West |
| WLR049 | R24 | | No | 59.906 | 10.764 | NIKU | Yes | Rostrum | Oslo | Norway | SL51844/F51873 | Star et al. 2018 | 1250 to 1350 | NA | NA | | | | | | West | Yes | 215841 | Yes | Yes | West | | West |
| WLR061 | R27 | | Yes | 59.907 | 10.763 | Museum of Cultural History | Yes | Rostrum | Oslo | Norway | C16818 KHM | Barrett et al. 2020 & this study | late 12th or 13th century based on Art History of zoomorphic carving | NA | NA | | | | | | East | Yes | 478005 | Yes | Yes | West | | West |
| WLR062 | R25 | | Yes | 59.906 | 10.768 | Museum of Cultural History | Yes | Rostrum | Oslo | Norway | C23798 KHM | Star et al. 2018 & this study | NA | NA | NA | | | | | | East | Yes | 50465653 | Yes | Yes | West | | West |
| WLR063 | FOC5374 | | No | 59.615 | 17.720 | Sigtuna Museum | Yes | Tusk (workshop debris) | Sigtuna | Sweden | kv Urmakaren; 8240a; Sigt ID 121830 | Star et al. 2018 | 1000 to 1050 | NA | NA | | | | | | East | Yes | 803098 | Yes | Yes | East | MT with Iceland specimens | East |
| WLR064 | FOC5375 | | No | 59.615 | 17.720 | Sigtuna Museum | Yes | Tusk (workshop debris) | Sigtuna | Sweden | kv Urmakaren; 8241a; Sigt ID 121831 | Star et al. 2018 | 1000 to 1025 | NA | NA | | | | | | East | Yes | 247469 | Yes | Yes | East | MT with Barents Sea specimens | East |
| WLR065 | FOC5376 | | Yes | 59.615 | 17.720 | Sigtuna Museum | Yes | Tusk (workshop debris) | Sigtuna | Sweden | kv Urmakaren; 8241b; Sigt ID 121831 | Star et al. 2018 & this study | 1000 to 1025 | NA | NA | | | | | | East | Yes | 62881481 | Yes | Yes | East | MT with Barents Sea specimens | East |
| WLR067 | FOC4848/R40 | | No | 59.616 | 17.719 | Sigtuna Museum | Yes | Rostrum | Sigtuna | Sweden | kv Trädgårdsmästaren; 29257 | Star et al. 2018 | 1200 to 1230 | NA | NA | | | | | | West | Yes | 954747 | Yes | Yes | West | | West |
| WLR068 | R51 | | No | 54.512 | 9.571 | Schleswig-Holsteinische Landesmuseen Schloss Gottorf | Yes | Rostrum | Schleswig | Germany | Schleswig Hafenstraße 13 | Star et al. 2018 | 12th to 13th century | NA | NA | | | | | | East | Yes | 1994450 | Yes | Yes | West | | West |
| WLR069 | WLR069 | | No | 60.987 | -45.423 | Natural History Museum of Denmark | Yes (within Greenland) | Rostrum | Igaliku/Gardar | Greenland | P9190912 P151/2017 KMG | Star et al. 2018 | Late 10th to 12th century | NA | NA | | | | | | East | Yes | 256034 | Yes | Yes | West | | West |
| WLR070 | WLR070 | | No | 60.987 | -45.423 | Natural History Museum of Denmark | Yes (within Greenland) | Rostrum | Igaliku/Gardar | Greenland | P9190871 P149/2017 KMG | Star et al. 2018 | Late 10th to 12th century | NA | NA | | | | | | East | Yes | 472034 | Yes | Yes | West | | West |
| WLR071 | WLR071 | | No | 60.987 | -45.423 | National Museum of Denmark | Yes (within Greenland) | Rostrum | Igaliku/Gardar | Greenland | PB091119 P153/2017 KMG | Star et al. 2018 | Late 10th to 12th century | NA | NA | | | | | | West | Yes | 235138 | Yes | Yes | West | | West |
| WLR072 | WLR072 | | No | 60.987 | -45.423 | National Museum of Denmark | Yes (within Greenland) | Rostrum | Igaliku/Gardar | Greenland | PB091098 P155/2017 KMG | Star et al. 2018 | Late 10th to 12th century | NA | NA | | | | | | West | Yes | 794178 | Yes | Yes | West | | West |
| WLR073 | R35 | | No | NA | NA | Musée Vert, muséum d'histoire naturelle du Mans | Yes | Rostrum | Le Mans (location of museum; find location uncertain) | France | MHNLM 2004.3.53 | Star et al. 2018 | 13th to 14th century | NA | NA | | | | | | West | Yes | 2594193 | Yes | Yes | West | | West |
| WLR074 | R64 | | No | 50.469 | 30.522 | Institute of Archaeology, National Academy of Sciences of Ukraine | Yes | Rostrum | Kyiv | Ukraine | 2319; sample S_07-2-2319 | Barrett et al. 2022 | late 12th century | NA | NA | | | | | | West | Yes | 742072 | Yes | Yes | West | | West |
| WLR075 | R65 | | No | 50.469 | 30.522 | Institute of Archaeology, National Academy of Sciences of Ukraine | Yes | Rostrum | Kyiv | Ukraine | 2369; sample S_07-1-2369 | Barrett et al. 2022 | late 12th century | NA | NA | | | | | | West | Yes | 247608 | Yes | Yes | West | | West |
| WLR076 | R66 | | No | 50.469 | 30.522 | Institute of Archaeology, National Academy of Sciences of Ukraine | Yes | Rostrum | Kyiv | Ukraine | 2825; sample S_07-1-2825 | Barrett et al. 2022 | late 12th century | NA | NA | | | | | | West | Yes | 451704 | Yes | Yes | West | | West |
| WLR077 | R71 | | Yes | 50.469 | 30.522 | Institute of Archaeology, National Academy of Sciences of Ukraine | Yes | Rostrum | Kyiv | Ukraine | 2937; sample S_11-2-2937 | Barrett et al. 2022 & this study | mid 12th century | NA | NA | | | | | | East | Yes | 154762865 | Yes | Yes | West | | West |
| WLR078 | R72 | | Yes | 50.469 | 30.522 | Institute of Archaeology, National Academy of Sciences of Ukraine | Yes | Rostrum | Kyiv | Ukraine | 2938; sample S_11-2-2938 | Barrett et al. 2022 & this study | mid 12th century | NA | NA | | | | | | East | Yes | 294070 | Yes | Yes | West | | West |
| WLR079 | R73 | | Yes | 50.469 | 30.522 | Institute of Archaeology, National Academy of Sciences of Ukraine | Yes | Rostrum | Kyiv | Ukraine | 2939; sample S_11-2-2939 | Barrett et al. 2022 & this study | mid 12th century | NA | NA | | | | | | West | Yes | 171145249 | Yes | Yes | West | | West |
| WLR080 | R74 | | No | 50.469 | 30.522 | Institute of Archaeology, National Academy of Sciences of Ukraine | Yes | Rostrum | Kyiv | Ukraine | 1833; sample S_11-3-1833 | Barrett et al. 2022 | mid 12th century | NA | NA | | | | | | West | Yes | 984168 | Yes | Yes | West | | West |
| WLR081 | FOC3018 | 25077 | NA | NA | NA | Natural History Museum London | ? | Other | Sápmi, northern Fennoscandia | Unknown | 1936.5.1.1 | This study | NA | Unknown | 1936 | 487 | 13 | 1884 | 1754 | 1938 | East | Yes | 3923032 | Yes | Yes | East | | East |
| WLR082 | FOC3061 | | NA | 59.616 | 17.720 | Sigtuna Museum | Yes | Tusk (workshop debris) | Sigtuna | Sweden | kv Trädgårdsmästaren 9-10; site find 30141; Sigt ID 0159608 | This study | 1175 to 1200 | NA | NA | | | | | | East | Yes | 173257 | Yes | Yes | West | | West |
| WLR083 | FOC3062 | | NA | 59.616 | 17.720 | Sigtuna Museum | Yes | Tusk (workshop debris) | Sigtuna | Sweden | kv Trädgårdsmästaren 9-10; site find 30141; Sigt ID 0159608 | This study | 1175 to 1200 | NA | NA | | | | | | East | Yes | 394231 | Yes | Yes | West | | West |
| WLR084 | FOC3063 | | NA | 59.616 | 17.720 | Sigtuna Museum | Yes | Tusk (workshop debris) | Sigtuna | Sweden | Professorn 1; site find 3460; Sigt ID 59521 | This study | 1130 to 1160 | NA | NA | | | | | | East | Yes | 457974 | Yes | Yes | East | MT with Iceland specimens | East |
| WLR085 | FOC3064 | | NA | 59.616 | 17.720 | Sigtuna Museum | Yes | Tusk (workshop debris) | Sigtuna | Sweden | Professorn 1; site find 4576; Sigt ID 60631 | This study | 1105 to 1130 | NA | NA | | | | | | West | Yes | 2927874 | Yes | Yes | West | | West |
| WLR086 | FOC3065 | | NA | 59.616 | 17.720 | Sigtuna Museum | Yes | Tusk (workshop debris) | Sigtuna | Sweden | Professorn 1; site find 9381; Sigt ID 65420 | This study | 1130 to 1160 | NA | NA | | | | | | West | Yes | 307591 | Yes | Yes | West | | West |
| WLR087 | FOC3066 | | NA | 59.616 | 17.720 | Sigtuna Museum | Yes | Rostrum (cheek tooth) | Sigtuna | Sweden | Professorn 1; site find 20643; Sigt ID 20832 | This study | 1160 to 1200 | NA | NA | | | | | | West | Yes | 240101 | Yes | Yes | West | | West |
| WLR088 | FOC3067 | | NA | 59.616 | 17.720 | Sigtuna Museum | Yes | Tusk (workshop debris) | Sigtuna | Sweden | Professorn 1; site find 46995; Sigt ID 102792 | This study | 1040 to 1055 | NA | NA | | | | | | East | Yes | 989553 | Yes | Yes | East | MT with Barents Sea specimens | East |
| WLR090 | FOC3069 | | NA | 59.616 | 17.720 | Sigtuna Museum | Yes | Tusk (workshop debris) | Sigtuna | Sweden | Professorn 1; site find 15236; Sigt ID 71269 | This study | 1130 to 1160 | NA | NA | | | | | | East | Yes | 1691563 | Yes | Yes | West | | West |
| WLR091 | FOC3071 | | NA | 59.616 | 17.720 | Sigtuna Museum | Yes | Tusk (broken object) | Sigtuna | Sweden | kv Trädgårdsmästaren 9-10; site find 958; Sigt ID 2174 | This study | 1200 to 1230 | NA | NA | | | | | | West | Yes | 323272 | Yes | Yes | West | | West |
| WLR092 | FOC3072 | | NA | 59.615 | 17.720 | Sigtuna Museum | Yes | Tusk (workshop debris) | Sigtuna | Sweden | Kv. Urmakaren 1; site find 8156; Sigt ID 38322 | This study | 1025 to 1050 | NA | NA | | | | | | East | Yes | 813961 | Yes | Yes | East | MT with Barents Sea specimens | East |
| WLR093 | FOC3073 | | NA | 59.615 | 17.720 | Sigtuna Museum | Yes | Tusk (workshop debris) | Sigtuna | Sweden | Kv. Urmakaren 1; site find 4881; Sigt ID 35050 | This study | 1000 to 1025 | NA | NA | | | | | | East | Yes | 146551 | Yes | Yes | East | MT with Iceland specimens | East |
| WLR094 | FOC3074/R41 | | NA | 59.616 | 17.719 | Sigtuna Museum | Yes | Rostrum | Sigtuna | Sweden | kv Trädgårdsmästaren; site find 23066; Sigt ID 23249 | This study | 1200 to 1230 | NA | NA | | | | | | East | Yes | 263851 | Yes | Yes | West | | West |
| WLR095 | FOC4327 | 21078 | NA | 63.476 | 7.757 | NTNU University Museum | Yes (but post-medieval) | Tusk (from shipwreck) | Skalmen fyr, Smøla | Norway | Aks. Nr. 2022/116 | This study | NA | NA | NA | 629 | 15 | 1803 | 1675 | 1953 | East | Yes | 1195634 | Yes | Yes | East | | East |
| WLR096 | FOC3828 | | No | 74.383 | -97.800 | Natural History Museum London | No | Rostrum | Barrow Strait | Canada | NHMUK ZD.1855.11.26.39 | Dierickx et al. 2025 | NA | 1854 | 1855 | | | | | | West | Yes | 3178570 | Yes | Yes | West | | West |
| WLR097 | FOC3833 | 22949 | No | 72.604 | -78.580 | Natural History Museum London | No | Other | "Pond's Bay, Davis Strait" (Mittimatalik/Pond Inlet?) | Canada | NHMUK ZD.1939.4474B | Dierickx et al. 2025 | NA | Unknown | 1939 | 604 | 13 | 1861 | 1720 | 1940 | East | Yes | 25534 | No | No | NA | MT with Iceland specimens | NA |
| WLR099 | FOC4616 | | No | NA | NA | Natural History Museum Oslo | No | Other | Greenland | Greenland | NHMO-DMA-32428/2-O | Dierickx et al. 2025 | NA | 1888 to 1889 inferred | 1893 | | | | | | East | Yes | 2442759 | Yes | Yes | West | | West |
| WLR100 | FOC4617 | | No | NA | NA | Natural History Museum Oslo | No | Other | Greenland | Greenland | NHMO-DMA-32428/4-O | Dierickx et al. 2025 | NA | 1888 to 1889 inferred | 1893 | | | | | | West | Yes | 7342215 | Yes | Yes | West | | West |
| WLR102 | FOC4669 | 22960 | No | NA | NA | Natural History Museum Oslo | No | Other | Greenland | Greenland | NHMO-DMA-31944 | Dierickx et al. 2025 | NA | 1844 | 1866 | | | | | | East | Yes | 5948197 | Yes | Yes | West | | West |
| WLR103 | FOC4819 | | No | NA | NA | Naturalis Biodiversity Center | No | Other | Greenland | Greenland | RMNH.MAM.64127 | Dierickx et al. 2025 | NA | Unknown | 1820 to 1863 | | | | | | * | Yes | 254308 | Yes | Yes | West | | West |
| WLR104 | FOC4845/R44 | 22954 | No | 52.094 | 5.118 | Naturalis Biodiversity Center | Yes | Rostrum | Utrecht | Netherlands | ZMA.MAM.7696 | Dierickx et al. 2025 | NA | NA | NA | 1325 | 16 | 1254 | 1127 | 1385 | East | Yes | 4513366 | Yes | Yes | West | | West |
| WLR105 | FOC3771 | 22937 | No | 47.360 | -61.880 | New Brunswick Museum | No | Other | Magdalen Islands | Canada | NBM-MA-001355 | Dierickx et al. 2025 | NA | NA | NA | 1210 | 16 | 1322 | 1103 | 1505 | West | Yes | 3630099 | Yes | Yes | West | | West |
| WLR106 | FOC4006 | 22424 | No | 47.493 | -61.755 | Private collection | No | Other | Magdalen Islands | Canada | NA | Dierickx et al. 2026 | NA | NA | NA | 683 | 22 | 1785 | 1583 | 1953 | West | Yes | 608636 | Yes | Yes | West | | West |
| WLR107 | FOC4332 | 23319 | No | 63.941 | -89.406 | Canadian Museum of History | No | Other | Silumuit | Canada | KkJg1:1490-DA01 | Dierickx et al. 2025 | NA | NA | NA | 868 | 13 | 1650 | 1510 | 1809 | West | Yes | 655142 | Yes | Yes | West | | West |
| WLR108 | FOC4419 | 24200 | No | 64.859 | -87.279 | Canadian Museum of History | No | Other | Kamarvik | Canada | LeHv-1:622.1-DA01 | Dierickx et al. 2025 | NA | NA | NA | 757 | 12 | 1774 | 1650 | 1940 | West | Yes | 964913 | Yes | Yes | West | | West |
| WLR109 | FOC4489 | | No | NA | NA | Canadian Museum of Nature | ? | Other | Unknown | Canada | 93DCA65 | Dierickx et al. 2025 | NA | NA | NA | | | | | | West | Yes | 1848099 | Yes | Yes | West | | West |
| WLR110 | FOC4496 | | No | 69.682 | -79.946 | Government of Nunavut Palaeontology Collection | No | Other | Jens Munk Island | Canada | NUFV 00439 | Dierickx et al. 2025 | NA | NA | NA | | | | | | West | Yes | 677116 | Yes | Yes | West | | West |
| WLR111 | FOC4510 | 24147 | No | 77.800 | -71.444 | Canadian Museum of Nature | No | Other | Neke, Cape Robertson | Greenland | CMNMA 9534 | Dierickx et al. 2025 | NA | Unknown | 1928 | 656 | 13 | 1817 | 1696 | 1927 | West | Yes | 5161770 | Yes | Yes | West | | West |
| WLR113 | FOC4538 | 24186 | No | 43.933 | -59.923 | Nova Scotia Museum | No | Other | Sable Island | Canada | NSM 78102 | Dierickx et al. 2025 | NA | NA | NA | 2601 | 17 | -246 | -427 | -32 | West | Yes | 538343 | Yes | Yes | West | | West |
| WLR114 | FOC4543 | 24190 | No | 43.933 | -59.923 | Sable Island Institute | No | Other | Sable Island | Canada | SII-00006 | Dierickx et al. 2025 | NA | NA | NA | 2052 | 14 | 424 | 241 | 606 | West | Yes | 651413 | Yes | Yes | West | | West |
| WLR115 | FOC4587 | 23521 | No | NA | NA | Svalbard Museum | No | Other | Svalbard | Norway | SVB 01010 | Dierickx et al. 2025 | NA | NA | NA | 538 | 12 | 1875 | 1729 | 1953 | * | Yes | 89617 | No | Yes | East* | Sufficient reads to confirm east, in combination with metadata | East |
| WLR116 | FOC4597 | 23505 | No | 78.833 | 28.000 | Tromsø Museum. The Arctic University Museum of Norway | No | Other | Kong Karls Land, Svalbard | Norway | TSZM 107 | Dierickx et al. 2025 | NA | NA | NA | 498 | 16 | 1892 | 1754 | 1953 | East | Yes | 4031328 | Yes | Yes | East | | East |
| WLR117 | FOC4600 | 23506 | No | 70.667 | 23.655 | Tromsø Museum. The Arctic University Museum of Norway | ? | Other | Hammerfest (Fuglenesset) | Norway | TSZM 117 | Dierickx et al. 2025 | NA | Unknown | 1929 | 575 | 12 | 1838 | 1703 | 1929 | East | Yes | 364744 | Yes | Yes | East | | East |
| WLR119 | FOC4619 | 23528 | No | NA | NA | Natural History Museum Oslo | No | Other | Svalbard | Norway | NHMO-DMA-33031 | Dierickx et al. 2025 | NA | NA | NA | 482 | 21 | 1895 | 1763 | 1953 | East | Yes | 1750884 | Yes | Yes | East | | East |
| WLR120 | FOC4663 | 23533 | No | 76.874 | 21.694 | Natural History Museum Oslo | No | Other | Svalbard (Edgeøya Håøya Tusenøyane) | Norway | NHMO-DMA-55942 | Dierickx et al. 2025 | NA | NA | NA | 478 | 13 | 1898 | 1770 | 1953 | East | Yes | 693754 | Yes | Yes | East | | East |
| WLR122 | FOC4668 | 23538 | No | 76.874 | 21.694 | Natural History Museum Oslo | No | Other | Svalbard (Edgeøya Håøya Tusenøyane) | Norway | NHMO-DMA-55964 | Dierickx et al. 2025 | NA | NA | NA | 560 | 14 | 1862 | 1717 | 1953 | East | Yes | 269465 | Yes | Yes | East | | East |
| WLR123 | FOC4836 | 24410 | No | 78.115 | 20.828 | Naturalis Biodiversity Center | No | Other | Kapp Lee area, Edgeøya, Svalbard | Norway | ZMA.MAM.16765 | Dierickx et al. 2025 | NA | NA | NA | 741 | 13 | 1687 | 1519 | 1856 | * | Yes | 80647 | No | Yes | East* | Sufficient reads to confirm east, in combination with metadata | East |
| WLR124 | FOC4841 | 23544 | No | 78.115 | 20.828 | Naturalis Biodiversity Center | No | Other | Kapp Lee, Edgeøya, Svalbard | Norway | ZMA.MAM.23411 | Dierickx et al. 2025 | NA | NA | NA | 789 | 14 | 1633 | 1489 | 1803 | East | Yes | 661942 | Yes | Yes | East | | East |
| WLR126 | FOC5377 | | NA | 59.615 | 17.719 | Sigtuna Museum | Yes | Tusk (crosier) | Sigtuna | Sweden | S:ta Gertrud 3 1993; site find 206; Sigt ID 159450 | This study | 11th to mid 13th century | NA | NA | | | | | | West | Yes | 2193866 | Yes | Yes | West | | West |

Table S2. Summary of aDNA results for the samples included in the combined mitochondrial and nuclear DNA analysis.

| Sample Name | ENA Accession | Raw Read Pairs (n) | PCR Duplicates | Uniquely Mapped Reads (n) | Endogenous DNA (fraction) | Reference Panel Specimen | Average Read Length | Nuclear coverage (fold) | X/mean autosome coverage ratio | Y/mean autosome coverage ratio | Y/X coverage ratio | Genetic sex (male/female) | Mapped Reads (MT) | MT Coverage (fold) | MT Clade | Power analyses | Sufficient reads for western assignment | Sufficient reads for eastern assignment | Nuclear Assignment Western Atlantic (prob) | Nuclear Assignment Eastern Atlantic (prob) | Nuclear SNPs (n) | Nuclear assignment | Plotted on mito-nuclear classification map (Figure 5) |
|---|---|---|---|---|---|---|---|---|---|---|---|---|---|---|---|---|---|---|---|---|---|---|---|
| WLR002 | ERS30746382 | 24811262 | 0.09 | 152320 | 0.01 | No | 89 | 0.00 | 0.9 | 0.0 | 0.0 | Female | 645 | 3.1 | West | No | Yes | Yes | 1 | 0 | 562 | West | Yes |
| WLR008 | ERS30746383 | 267454144 | 0.03 | 138420421 | 0.54 | Yes | 75 | 4.07 | 0.5 | 0.7 | 1.4 | Male | 255765 | 1058 | East | Yes | Yes | Yes | 0 | 1 | 142464 | East | Yes |
| WLR009 | ERS30746384 | 16314474 | 0.02 | 1513126 | 0.09 | No | 76 | 0.05 | 0.5 | 0.6 | 1.7 | Male | 3919 | 17 | East | Yes | Yes | Yes | 0 | 1 | 8109 | East | Yes |
| WLR010 | ERS30746385 | 19358379 | 0.01 | 6531818 | 0.33 | Yes | 83 | 0.22 | 0.5 | 0.8 | 1.6 | Male | 13755 | 61 | East | Yes | Yes | Yes | 0 | 1 | 33301 | East | Yes |
| WLR011 | ERS30746386 | 6182878 | 0.00 | 1466898 | 0.24 | No | 67 | 0.04 | 0.9 | 0.0 | 0.0 | Female | 2542 | 9 | East | Yes | Yes | Yes | 0 | 1 | 6019 | East | Yes |
| WLR012 | ERS30746387 | 60602845 | 0.01 | 35780426 | 0.61 | Yes | 74 | 1.08 | 0.5 | 0.7 | 1.4 | Male | 103517 | 424 | East | Yes | Yes | Yes | 0 | 1 | 103457 | East | Yes |
| WLR017 | ERS30746388 | 245060010 | 0.02 | 142933996 | 0.60 | Yes | 75 | 4.24 | 0.9 | 0.0 | 0.0 | Female | 316011 | 1345 | East | Yes | Yes | Yes | 0 | 1 | 142735 | East | Yes |
| WLR021 | ERS30746389 | 232662312 | 0.06 | 150582649 | 0.65 | Yes | 96 | 5.36 | 0.5 | 0.9 | 1.6 | Male | 245678 | 1240 | East | Yes | Yes | Yes | 0 | 1 | 143941 | East | Yes |
| WLR022 | ERS30746390 | 159179202 | 0.04 | 93360443 | 0.59 | Yes | 96 | 3.26 | 0.5 | 0.8 | 1.5 | Male | 180079 | 885 | East | Yes | Yes | Yes | 0 | 1 | 140329 | East | Yes |
| WLR023 | ERS30746391 | 11769650 | 0.01 | 4338170 | 0.37 | Yes | 95 | 0.18 | 0.5 | 0.8 | 1.8 | Male | 14682 | 77 | East | Yes | Yes | Yes | 0 | 1 | 27632 | East | Yes |
| WLR024 | ERS30746392 | 214389942 | 0.05 | 139197896 | 0.65 | Yes | 94 | 4.69 | 0.5 | 0.8 | 1.5 | Male | 155786 | 745 | East | Yes | Yes | Yes | 0 | 1 | 143503 | East | Yes |
| WLR029 | ERS30746393 | 11530949 | 0.02 | 3669169 | 0.31 | Yes | 84 | 0.11 | 0.5 | 0.6 | 1.4 | Male | 13177 | 61 | East | Yes | Yes | Yes | 1 | 0 | 17306 | West | Yes |
| WLR030 | ERS30746394 | 16913487 | 0.03 | 3320527 | 0.19 | Yes | 82 | 0.10 | 0.5 | 0.6 | 1.5 | Male | 20673 | 98 | West | Yes | Yes | Yes | 1 | 0 | 16208 | West | Yes |
| WLR031 | ERS30746395 | 9453976 | 0.02 | 1920148 | 0.19 | No | 90 | 0.06 | 0.5 | 0.5 | 1.5 | Male | 25374 | 123 | West | Yes | Yes | Yes | 1 | 0 | 9266 | West | Yes |
| WLR033 | ERS30746396 | 17401591 | 0.04 | 210538 | 0.01 | No | 105 | 0.01 | 0.5 | 0.2 | 1.6 | Male | 1485 | 9 | West | No | Yes | Yes | 1 | 0 | 1305 | West | Yes |
| WLR034 | ERS30746397 | 30144644 | 0.42 | 598889 | 0.02 | No | 76 | 0.01 | 0.5 | 0.2 | 1.5 | Male | 1795 | 7 | East | Yes | Yes | Yes | 1 | 0 | 2321 | West | Yes |
| WLR038 | ERS30746398 | 398716111 | 0.05 | 27988033 | 0.08 | Yes | 84 | 0.83 | 0.5 | 0.8 | 1.6 | Male | 69245 | 326 | East | Yes | Yes | Yes | 1 | 0 | 89538 | West | Yes |
| WLR039 | ERS30746399 | 8490526 | 0.03 | 87152 | 0.01 | No | 102 | 0.00 | 0.9 | 0.0 | 0.1 | Female | 2438 | 14 | West | No | No | Yes | NA | 0 | 339 | NA | No |
| WLR040 | ERS30746400 | 117667362 | 0.41 | 102003 | 0.00 | No | 94 | 0.00 | 0.7 | 0.1 | 1.7 | Male | 1967 | 9 | East | No | No | Yes | NA | 0.70 | 62 | NA | No |
| WLR041 | ERS30746401 | 393024631 | 0.06 | 61352584 | 0.16 | Yes | 85 | 1.77 | 0.5 | 0.8 | 1.6 | Male | 251592 | 1102 | West | Yes | Yes | Yes | 1 | 0 | 125195 | West | Yes |
| WLR042 | ERS30746402 | 11013196 | 0.03 | 141584 | 0.01 | No | 123 | 0.00 | 0.5 | 0.1 | 1.6 | Male | 2264 | 15 | West | No | Yes | Yes | 1 | 0 | 693 | West | Yes |
| WLR043 | ERS30746403 | 13253610 | 0.10 | 80994 | 0.01 | No | 98 | 0.00 | 0.8 | 0.0 | 0.9 | Male | 1755 | 9 | East | No | No | Yes | NA | 0 | 323 | NA | No |
| WLR044 | ERS30746404 | 253370015 | 0.06 | 94591655 | 0.38 | Yes | 91 | 3.02 | 0.5 | 0.9 | 1.6 | Male | 72541 | 386 | West | Yes | Yes | Yes | 1 | 0 | 139275 | West | Yes |
| WLR045 | ERS30746405 | 11964390 | 0.03 | 1430046 | 0.12 | No | 83 | 0.05 | 1.0 | 0.0 | 0.0 | Female | 5285 | 26 | West | Yes | Yes | Yes | 1 | 0 | 6831 | West | Yes |
| WLR046 | ERS30746406 | 9666872 | 0.03 | 283791 | 0.03 | No | 109 | 0.01 | 0.5 | 0.3 | 2.1 | Male | 3670 | 25 | West | No | Yes | Yes | 1 | 0 | 1754 | West | Yes |
| WLR047 | ERS30746407 | 11236317 | 0.05 | 3420742 | 0.30 | No | 74 | 0.09 | 0.5 | 0.6 | 1.6 | Male | 2424 | 9 | East | Yes | Yes | Yes | 1 | 0 | 14103 | West | Yes |
| WLR048 | ERS30746408 | 281637982 | 0.05 | 18113210 | 0.06 | Yes | 134 | 0.60 | 0.5 | 0.8 | 1.5 | Male | 166975 | 899 | West | Yes | Yes | Yes | 1 | 0 | 72247 | West | Yes |
| WLR049 | ERS30746409 | 12337684 | 0.03 | 215841 | 0.02 | No | 108 | 0.01 | 1.0 | 0.0 | 0.0 | Female | 1724 | 10 | West | No | Yes | Yes | 1 | 0 | 1092 | West | Yes |
| WLR061 | ERS30746410 | 184581406 | 0.37 | 478005 | 0.00 | No | 68 | 0.01 | 0.7 | 0.0 | 0.0 | Female | 2788 | 11 | East | No | Yes | Yes | 1 | 0 | 1228 | West | Yes |
| WLR062 | ERS30746411 | 283571273 | 0.09 | 50465653 | 0.26 | Yes | 68 | 1.32 | 0.5 | 0.8 | 1.5 | Male | 528238 | 2074 | East | Yes | Yes | Yes | 1 | 0 | 112479 | West | Yes |
| WLR063 | ERS30746412 | 12129433 | 0.01 | 803098 | 0.07 | No | 95 | 0.03 | 0.6 | 0.3 | 1.4 | Male | 10398 | 52 | East | Yes | Yes | Yes | 0 | 1 | 3976 | East | Yes |
| WLR064 | ERS30746413 | 8742672 | 0.01 | 247469 | 0.03 | No | 101 | 0.01 | 0.5 | 0.1 | 1.1 | Male | 2962 | 14 | East | No | Yes | Yes | 0 | 1 | 1181 | East | Yes |
| WLR065 | ERS30746414 | 323614870 | 0.06 | 62881481 | 0.21 | No | 88 | 1.77 | 0.5 | 0.7 | 1.3 | Male | 1235739 | 5818 | East | Yes | Yes | Yes | 0 | 1 | 119998 | East | Yes |
| WLR067 | ERS30746415 | 11302823 | 0.02 | 954747 | 0.08 | No | 84 | 0.03 | 0.5 | 0.4 | 1.5 | Male | 6134 | 30 | West | Yes | Yes | Yes | 1 | 0 | 4505 | West | Yes |
| WLR068 | ERS30746416 | 10035185 | 0.02 | 1994450 | 0.20 | No | 78 | 0.06 | 0.5 | 0.5 | 1.6 | Male | 26220 | 112 | East | Yes | Yes | Yes | 1 | 0 | 8695 | West | Yes |
| WLR069 | ERS30746417 | 23155468 | 0.01 | 256034 | 0.01 | No | 72 | 0.01 | 1.0 | 0.0 | 0.0 | Female | 1187 | 5 | East | No | Yes | Yes | 1 | 0 | 1257 | West | Yes |
| WLR070 | ERS30746418 | 27296103 | 0.01 | 472034 | 0.02 | No | 75 | 0.01 | 1.0 | 0.0 | 0.0 | Female | 2591 | 12 | East | No | Yes | Yes | 1 | 0 | 2236 | West | Yes |
| WLR071 | ERS30746419 | 27049413 | 0.02 | 235138 | 0.01 | No | 73 | 0.01 | 0.5 | 0.1 | 1.5 | Male | 1457 | 6 | West | No | Yes | Yes | 1 | 0 | 1138 | West | Yes |
| WLR072 | ERS30746420 | 34627011 | 0.01 | 794178 | 0.02 | No | 73 | 0.02 | 0.5 | 0.3 | 1.3 | Male | 2519 | 11 | West | Yes | Yes | Yes | 1 | 0 | 3724 | West | Yes |
| WLR073 | ERS30746421 | 16056610 | 0.02 | 2594193 | 0.16 | No | 71 | 0.08 | 1.0 | 0.0 | 0.0 | Female | 15420 | 74 | West | Yes | Yes | Yes | 1 | 0 | 11593 | West | Yes |
| WLR074 | ERS30746422 | 5399576 | 0.20 | 742072 | 0.14 | No | 52 | 0.02 | 0.5 | 0.2 | 1.4 | Male | 5759 | 19 | West | Yes | Yes | Yes | 1 | 0 | 2694 | West | Yes |
| WLR075 | ERS30746423 | 5719053 | 0.21 | 247608 | 0.04 | No | 67 | 0.01 | 0.5 | 0.1 | 1.7 | Male | 1375 | 5 | West | No | Yes | Yes | 1 | 0 | 975 | West | Yes |
| WLR076 | ERS30746424 | 7891733 | 0.21 | 451704 | 0.06 | No | 62 | 0.01 | 0.5 | 0.2 | 1.5 | Male | 2761 | 11 | West | No | Yes | Yes | 1 | 0 | 1853 | West | Yes |
| WLR077 | ERS30746425 | 416842471 | 0.07 | 154762865 | 0.38 | Yes | 77 | 4.34 | 0.5 | 0.7 | 1.4 | Male | 305156 | 1290 | East | Yes | Yes | Yes | 1 | 0 | 143405 | West | Yes |
| WLR078 | ERS30746426 | 13485498 | 0.33 | 294070 | 0.02 | No | 58 | 0.01 | 0.8 | 0.0 | 0.0 | Female | 1592 | 5 | East | No | Yes | Yes | 1 | 0 | 897 | West | Yes |
| WLR079 | ERS30746427 | 933262111 | 0.10 | 171145249 | 0.20 | Yes | 79 | 4.71 | 0.9 | 0.0 | 0.0 | Female | 564219 | 2483 | West | Yes | Yes | Yes | 1 | 0 | 143322 | West | Yes |
| WLR080 | ERS30746428 | 5492488 | 0.18 | 984168 | 0.18 | No | 67 | 0.02 | 0.5 | 0.3 | 1.3 | Male | 2994 | 11 | West | Yes | Yes | Yes | 1 | 0 | 3840 | West | Yes |
| WLR081 | ERS30746429 | 13262146 | 0.07 | 3923032 | 0.30 | No | 77 | 0.12 | 0.5 | 0.9 | 2.3 | Male | 3700 | 16 | East | Yes | Yes | Yes | 0 | 1 | 17175 | East | Yes |
| WLR082 | ERS30746430 | 12916374 | 0.09 | 173257 | 0.01 | No | 118 | 0.00 | 0.5 | 0.1 | 1.4 | Male | 1270 | 5 | East | No | Yes | Yes | 1 | 0 | 666 | West | Yes |
| WLR083 | ERS30746431 | 24735249 | 0.06 | 394231 | 0.02 | No | 94 | 0.01 | 0.5 | 0.1 | 1.2 | Male | 32322 | 126 | East | No | Yes | Yes | 1 | 0 | 1385 | West | Yes |
| WLR084 | ERS30746432 | 43370798 | 0.07 | 457974 | 0.01 | No | 95 | 0.01 | 0.5 | 0.2 | 1.2 | Male | 2691 | 11 | East | No | Yes | Yes | 0 | 1 | 1725 | East | Yes |
| WLR085 | ERS30746433 | 8436802 | 0.07 | 2927874 | 0.38 | No | 83 | 0.08 | 1.0 | 0.0 | 0.0 | Female | 43682 | 209 | West | Yes | Yes | Yes | 1 | 0 | 12126 | West | Yes |
| WLR086 | ERS30746434 | 6516418 | 0.07 | 307591 | 0.05 | No | 82 | 0.01 | 1.0 | 0.0 | 0.0 | Female | 7557 | 29 | West | No | Yes | Yes | 1 | 0 | 1088 | West | Yes |
| WLR087 | ERS30746435 | 6576366 | 0.10 | 240101 | 0.04 | No | 93 | 0.01 | 1.0 | 0.0 | 0.0 | Female | 7524 | 31 | West | No | Yes | Yes | 1 | 0 | 1040 | West | Yes |
| WLR088 | ERS30746436 | 8702427 | 0.07 | 989553 | 0.13 | No | 84 | 0.03 | 0.5 | 0.3 | 1.3 | Male | 12979 | 60 | East | Yes | Yes | Yes | 0 | 1 | 4609 | East | Yes |
| WLR090 | ERS30746437 | 5391944 | 0.06 | 1691563 | 0.33 | No | 74 | 0.05 | 0.5 | 0.5 | 1.6 | Male | 8025 | 32 | East | Yes | Yes | Yes | 1 | 0 | 7750 | West | Yes |
| WLR091 | ERS30746438 | 12416692 | 0.12 | 323272 | 0.05 | No | 79 | 0.01 | 0.9 | 0.0 | 0.0 | Female | 14103 | 46 | West | No | Yes | Yes | 1 | 0 | 869 | West | Yes |
| WLR092 | ERS30746439 | 66944955 | 0.11 | 813961 | 0.01 | No | 82 | 0.02 | 0.5 | 0.2 | 1.3 | Male | 15220 | 56 | East | Yes | Yes | Yes | 0 | 1 | 2695 | East | Yes |
| WLR093 | ERS30746440 | 8546957 | 0.06 | 146551 | 0.02 | No | 95 | 0.00 | 0.5 | 0.1 | 1.5 | Male | 1618 | 8 | East | No | Yes | Yes | 0 | 1 | 653 | East | Yes |
| WLR094 | ERS30746441 | 8597369 | 0.03 | 263851 | 0.04 | No | 86 | 0.01 | 0.9 | 0.0 | 0.0 | Female | 1043 | 4 | East | No | Yes | Yes | 1 | 0 | 1039 | West | Yes |
| WLR095 | ERS30746442 | 4039508 | 0.02 | 1195634 | 0.58 | No | 95 | 0.05 | 0.5 | 0.5 | 1.6 | Male | 9907 | 52 | East | Yes | Yes | Yes | 0 | 1 | 7167 | East | No |
| WLR096 | ERS30746443 | 6282935 | 0.02 | 3178570 | 0.53 | No | 60 | 0.08 | 0.5 | 0.8 | 2.2 | Male | 9367 | 33 | West | Yes | Yes | Yes | 1 | 0 | 11255 | West | Yes |
| WLR097 | ERS30746444 | 12611169 | 0.44 | 25534 | 0.00 | No | 86 | 0.00 | 0.4 | 0.0 | 0.9 | Male | 691 | 3 | East | No | No | No | NA | NA | 50 | NA | No |
| WLR099 | ERS30746445 | 7837980 | 0.02 | 2442759 | 0.37 | No | 64 | 0.06 | 1.0 | 0.0 | 0.0 | Female | 5984 | 22 | East | Yes | Yes | Yes | 1 | 0 | 9384 | West | Yes |
| WLR100 | ERS30746446 | 12594983 | 0.03 | 7342215 | 0.61 | No | 61 | 0.18 | 1.0 | 0.0 | 0.0 | Female | 5969 | 21 | West | Yes | Yes | Yes | 1 | 0 | 25595 | West | Yes |
| WLR102 | ERS30746447 | 8881104 | 0.02 | 5948197 | 0.70 | No | 64 | 0.16 | 0.5 | 0.7 | 1.5 | Male | 4425 | 17 | East | Yes | Yes | Yes | 1 | 0 | 22934 | West | Yes |
| WLR103 | ERS30746448 | 16739426 | 0.06 | 254308 | 0.02 | No | 63 | 0.01 | 1.0 | 0.0 | 0.0 | Female | 514 | 2 | NA | No | Yes | Yes | 1 | 0 | 876 | West | Yes |
| WLR104 | ERS30746449 | 9951847 | 0.03 | 4513366 | 0.50 | No | 63 | 0.11 | 0.5 | 0.6 | 1.5 | Male | 4766 | 18 | East | Yes | Yes | Yes | 1 | 0 | 16755 | West | Yes |
| WLR105 | ERS30746450 | 13716199 | 0.13 | 3630099 | 0.35 | No | 79 | 0.09 | 1.0 | 0.0 | 0.0 | Female | 13412 | 55 | West | Yes | Yes | Yes | 1 | 0 | 13794 | West | Yes |
| WLR106 | ERS30746451 | 9619538 | 0.03 | 608636 | 0.07 | No | 67 | 0.02 | 0.5 | 0.3 | 1.4 | Male | 1017 | 4 | West | Yes | Yes | Yes | 1 | 0 | 2684 | West | Yes |
| WLR107 | ERS30746452 | 13826944 | 0.04 | 655142 | 0.05 | No | 65 | 0.01 | 0.5 | 0.2 | 1.5 | Male | 1233 | 4 | West | Yes | Yes | Yes | 1 | 0 | 2190 | West | Yes |
| WLR108 | ERS30746453 | 5227622 | 0.02 | 964913 | 0.19 | No | 84 | 0.04 | 0.5 | 0.4 | 1.6 | Male | 3006 | 16 | West | Yes | Yes | Yes | 1 | 0 | 5654 | West | Yes |

| Sample Name | ENA Accession | Raw Read Pairs (n) | PCR Duplicates | Uniquely Mapped Reads (n) | Endogenous DNA (fraction) | Reference Panel Specimen | Average Read Length | Nuclear coverage (fold) | X/mean autosome coverage ratio | Y/mean autosome coverage ratio | Y/X coverage ratio | Genetic sex (male/female) | Mapped Reads (MT) | MT Coverage (fold) | MT Clade | Power analyses | Sufficient reads for western assignment | Sufficient reads for eastern assignment | Nuclear Assignment Western Atlantic (prob) | Nuclear Assignment Eastern Atlantic (prob) | Nuclear SNPs (n) | Nuclear assignment | Plotted on mito-nuclear classification map (Figure 5) |
|---|---|---|---|---|---|---|---|---|---|---|---|---|---|---|---|---|---|---|---|---|---|---|---|
| WLR109 | ERS30746454 | 9212273 | 0.06 | 1848099 | 0.25 | No | 83 | 0.05 | 0.5 | 0.5 | 1.5 | Male | 7057 | 28 | West | Yes | Yes | Yes | 1 | 0 | 7613 | West | Yes |
| WLR110 | ERS30746455 | 5172564 | 0.05 | 677116 | 0.14 | No | 88 | 0.02 | 0.5 | 0.3 | 1.5 | Male | 2004 | 8 | West | Yes | Yes | Yes | 1 | 0 | 3506 | West | Yes |
| WLR111 | ERS30746456 | 10015385 | 0.02 | 5161770 | 0.54 | No | 68 | 0.14 | 1.1 | 0.0 | 0.0 | Female | 7355 | 25 | West | Yes | Yes | Yes | 1 | 0 | 19323 | West | Yes |
| WLR113 | ERS30746457 | 6509399 | 0.05 | 538343 | 0.09 | No | 86 | 0.02 | 0.5 | 0.2 | 1.5 | Male | 2131 | 10 | West | Yes | Yes | Yes | 1 | 0 | 2398 | West | Yes |
| WLR114 | ERS30746458 | 7701363 | 0.13 | 651413 | 0.10 | No | 78 | 0.02 | 0.5 | 0.3 | 1.5 | Male | 1870 | 8 | West | Yes | Yes | Yes | 1 | 0 | 2742 | West | Yes |
| WLR115 | ERS30746459 | 4142652 | 0.08 | 89617 | 0.02 | No | 65 | 0.00 | 0.5 | 0.0 | 1.4 | Male | 319 | 1 | NA | No | No | Yes | NA | 1 | 283 | East* | Yes |
| WLR116 | ERS30746460 | 5963428 | 0.02 | 4031328 | 0.69 | No | 80 | 0.13 | 1.0 | 0.0 | 0.0 | Female | 4878 | 22 | East | Yes | Yes | Yes | 0 | 1 | 18443 | East | Yes |
| WLR117 | ERS30746461 | 8322178 | 0.03 | 364744 | 0.05 | No | 87 | 0.01 | 0.5 | 0.2 | 1.4 | Male | 2025 | 8 | East | No | Yes | Yes | 0 | 1 | 1539 | East | Yes |
| WLR119 | ERS30746462 | 5406174 | 0.06 | 1750884 | 0.42 | No | 69 | 0.05 | 0.5 | 0.5 | 1.5 | Male | 41041 | 152 | East | Yes | Yes | Yes | 0 | 1 | 6742 | East | Yes |
| WLR120 | ERS30746463 | 5159365 | 0.04 | 693754 | 0.15 | No | 72 | 0.02 | 0.5 | 0.3 | 1.3 | Male | 4367 | 17 | East | Yes | Yes | Yes | 0 | 1 | 3020 | East | Yes |
| WLR122 | ERS30746464 | 8124487 | 0.05 | 269465 | 0.04 | No | 82 | 0.01 | 1.0 | 0.0 | 0.0 | Female | 1461 | 6 | East | No | Yes | Yes | 0 | 1 | 1295 | East | Yes |
| WLR123 | ERS30746465 | 6895842 | 0.21 | 80647 | 0.01 | No | 77 | 0.00 | 0.8 | 0.0 | 0.0 | Female | 588 | 2 | NA | No | No | Yes | NA | 1 | 277 | East* | Yes |
| WLR124 | ERS30746466 | 6941922 | 0.05 | 661942 | 0.11 | No | 83 | 0.02 | 1.0 | 0.0 | 0.0 | Female | 10151 | 44 | East | Yes | Yes | Yes | 0 | 1 | 3081 | East | Yes |
| WLR126 | ERS30746467 | 15494733 | 0.16 | 2193866 | 0.25 | No | 83 | 0.06 | 0.5 | 0.5 | 1.4 | Male | 26572 | 103 | West | Yes | Yes | Yes | 1 | 0 | 8951 | West | Yes |

Table S3. Sample details for walrus specimens with mitogenomes published by Keighley et al. (2019) and Ruiz-Puerta et al. (2023) that have been employed as comparative control data.

| aDNA Unique Identifier | Country | Original Identifier | Data First Published | MT Clade | Available Nuclear Data |
|---|---|---|---|---|---|
| MK671108 | Greenland | XOR#017 ZM#293 | Keighley et al. 2019 | East | No |
| MK671123 | Canada | XOR#061 TEAL-6593 | Keighley et al. 2019 | West | No |
| MK671128 | Iceland | XOR#031 | Keighley et al. 2019 | East | No |
| MK671124 | Iceland | XOR#112 | Keighley et al. 2019 | East | No |
| ER0112 | Canada | CMNFV 35797 | Ruiz-Puerta et al. 2023 | West | No |
| ER0113 | Canada | CMNFV 43818 | Ruiz-Puerta et al. 2023 | West | No |
| ER0114 | Canada | CMNFV 43819 | Ruiz-Puerta et al. 2023 | West | No |
| ER0116 | Canada | CMNFV 43821 | Ruiz-Puerta et al. 2023 | West | No |
| ER0117 | Canada | CMNFV 43825 | Ruiz-Puerta et al. 2023 | West | No |
| ER0118 | Canada | CMNFV 48378 | Ruiz-Puerta et al. 2023 | East | No |
| ER0119 | Canada | CMNFV 48389 | Ruiz-Puerta et al. 2023 | East | No |
| ER0120 | Canada | CMNFV 48390 | Ruiz-Puerta et al. 2023 | East | No |
| ER0121 | Canada | CMNFV 55070 | Ruiz-Puerta et al. 2023 | East | No |
| ER0122 | Canada | CMNFV 55135 | Ruiz-Puerta et al. 2023 | East | No |
| ER0124 | Canada | CMNFV 56803 | Ruiz-Puerta et al. 2023 | East | No |
| ER0202 | Canada | NgHd-1:01624 | Ruiz-Puerta et al. 2023 | East | No |
| ER0205 | Canada | NgHd-1:02806 | Ruiz-Puerta et al. 2023 | West | No |
| ER0206 | Canada | NgHd-1:04364 | Ruiz-Puerta et al. 2023 | West | No |
| ER0207 | Canada | NgHd-1:04432 | Ruiz-Puerta et al. 2023 | West | No |
| ER0208 | Canada | NgHd-1:05872 | Ruiz-Puerta et al. 2023 | West | No |
| ER0209 | Canada | NgHd-1:06299 | Ruiz-Puerta et al. 2023 | East | No |
| ER0211 | Canada | NgHd-1:07866 | Ruiz-Puerta et al. 2023 | East | No |
| ER0212 | Canada | NgHd-1:07906 | Ruiz-Puerta et al. 2023 | West | No |
| ER0213 | Canada | NgHd-1:10039 | Ruiz-Puerta et al. 2023 | West | No |
| ER0216 | Canada | NgHd-1:10668 | Ruiz-Puerta et al. 2023 | East | No |
| ER0221 | Canada | NUFV 456 | Ruiz-Puerta et al. 2023 | East | No |
| XOR001 | Greenland | INWsample059 | Ruiz-Puerta et al. 2023 | West | No |
| XOR002 | Greenland | INWsample029 | Ruiz-Puerta et al. 2023 | West | No |
| XOR004 | Greenland | INWsample018 | Ruiz-Puerta et al. 2023 | West | No |
| XOR008 | Greenland | M1269 | Ruiz-Puerta et al. 2023 | West | No |
| XOR009 | Greenland | 403 | Ruiz-Puerta et al. 2023 | West | No |
| XOR018 | Greenland | INWsample066 | Ruiz-Puerta et al. 2023 | West | No |
| XOR020 | Greenland | CN461 | Ruiz-Puerta et al. 2023 | East | No |
| XOR022 | Greenland | ZMK73/1997 | Ruiz-Puerta et al. 2023 | East | No |
| XOR023 | Greenland | 465 | Ruiz-Puerta et al. 2023 | East | No |
| XOR028 | Greenland | 385 | Ruiz-Puerta et al. 2023 | East | No |
| XOR035 | Greenland | ZMK136/1989 | Ruiz-Puerta et al. 2023 | West | No |

| aDNA Unique Identifier | Country | Original Identifier | Data First Published | MT Clade | Available Nuclear Data |
|---|---|---|---|---|---|
| XOR043 | Greenland | ZMK135/1989 | Ruiz-Puerta et al. 2023 | East | No |
| XOR044 | Greenland | ZMK3/1978 | Ruiz-Puerta et al. 2023 | West | No |
| XOR053 | Canada | TEAL-6328 | Ruiz-Puerta et al. 2023 | East | No |
| XOR056 | Canada | TEAL-6598 | Ruiz-Puerta et al. 2023 | West | No |
| XOR060 | Canada | TEAL-7523 | Ruiz-Puerta et al. 2023 | East | No |
| XOR065 | Canada | TEAL-7549 | Ruiz-Puerta et al. 2023 | West | No |
| XOR080 | Greenland | ZMK136/1989 | Ruiz-Puerta et al. 2023 | West | No |
| XOR081 | Greenland | ZMK136/1989 | Ruiz-Puerta et al. 2023 | East | No |
| XOR082 | Greenland | ZMK136/1989 | Ruiz-Puerta et al. 2023 | East | No |
| XOR086 | Greenland | ZMK73/1997 | Ruiz-Puerta et al. 2023 | West | No |
| XOR087 | Greenland | ZMK65/2008 | Ruiz-Puerta et al. 2023 | East | No |
| XOR091 | Greenland | ZMK68/1996 | Ruiz-Puerta et al. 2023 | West | No |
| XOR094 | Greenland | ZMK126/2007 | Ruiz-Puerta et al. 2023 | East | No |
| XOR097 | Iceland | MOR#16 | Ruiz-Puerta et al. 2023 | East | No |
| XOR098 | Iceland | MOR#11 | Ruiz-Puerta et al. 2023 | East | No |
| XOR099 | Iceland | MOR#29 | Ruiz-Puerta et al. 2023 | East | No |
| XOR111 | Iceland | MOR#18 | Ruiz-Puerta et al. 2023 | East | No |
| XOR113 | Iceland | MOR# 25 | Ruiz-Puerta et al. 2023 | East | No |
| XOR114 | Iceland | MOR#26 | Ruiz-Puerta et al. 2023 | East | No |
| XOR118 | Greenland | ZMK68/1996 | Ruiz-Puerta et al. 2023 | West | No |
| XOR119 | Greenland | ZMK63/2008 | Ruiz-Puerta et al. 2023 | East | No |
| XOR120 | Canada | NhHd-3:3948 | Ruiz-Puerta et al. 2023 | West | No |
| XOR123 | Canada | NhHd-7:205 | Ruiz-Puerta et al. 2023 | West | No |
| XOR125 | Canada | NhHd-9:713 | Ruiz-Puerta et al. 2023 | West | No |
| XOR126 | Canada | NdHd-9:634 | Ruiz-Puerta et al. 2023 | West | No |
| XOR127 | Greenland | ZMK 54/1993 | Ruiz-Puerta et al. 2023 | East | No |
| XOR130 | Greenland | ZMK50/1993 | Ruiz-Puerta et al. 2023 | East | No |
| XOR135 | Canada | ZMK123a/1955 | Ruiz-Puerta et al. 2023 | East | No |
| XOR136 | Canada | ZMK123a/1955 | Ruiz-Puerta et al. 2023 | West | No |
| XOR139 | Greenland | ZMK50/1993 | Ruiz-Puerta et al. 2023 | East | No |
| XOR141 | Greenland | ZMK119/2007 | Ruiz-Puerta et al. 2023 | East | No |
| XOR146 | Canada | ZMK123a/1955 | Ruiz-Puerta et al. 2023 | West | No |
| XOR147 | Canada | ZMK123a/1955 | Ruiz-Puerta et al. 2023 | West | No |
| XOR151 | Greenland | ZMK71/1997 | Ruiz-Puerta et al. 2023 | West | No |
| XOR153 | Greenland | ZMK19/1933 | Ruiz-Puerta et al. 2023 | East | No |
| XOR155 | Canada | ZMK123a/1955 | Ruiz-Puerta et al. 2023 | East | No |
| XOR157 | Canada | ZMK123a/1955 | Ruiz-Puerta et al. 2023 | East | No |
| XOR158 | Canada | ZMK123c/1955 | Ruiz-Puerta et al. 2023 | East | No |

| aDNA Unique Identifier | Country | Original Identifier | Data First Published | MT Clade | Available Nuclear Data |
|---|---|---|---|---|---|
| XOR159 | Canada | ZMK123c/1955 | Ruiz-Puerta et al. 2023 | West | No |
| XOR160 | Iceland | MOR#22 | Ruiz-Puerta et al. 2023 | East | No |
| XOR161 | Iceland | MOR#17 | Ruiz-Puerta et al. 2023 | East | No |
| XOR162 | Iceland | MOR#23 | Ruiz-Puerta et al. 2023 | East | No |
| XOR163 | Iceland | MOR#14 | Ruiz-Puerta et al. 2023 | East | No |
| XOR164 | Iceland | MOR#32 | Ruiz-Puerta et al. 2023 | East | No |
| XOR165 | Canada | NhHd-7:349 | Ruiz-Puerta et al. 2023 | East | No |
| XOR166 | Canada | NhHd-9:264 | Ruiz-Puerta et al. 2023 | West | No |
| XOR167 | Canada | NhHd-3:1454 | Ruiz-Puerta et al. 2023 | West | No |
| XOR168 | Canada | NhHd-1:DNA1 | Ruiz-Puerta et al. 2023 | West | No |
| XOR169 | Canada | NhHd-1:DNA2 | Ruiz-Puerta et al. 2023 | West | No |
| XOR172 | Canada | NhHe-11:DNA5 | Ruiz-Puerta et al. 2023 | West | No |
| XOR173 | Canada | NhHe-11:DNA6 | Ruiz-Puerta et al. 2023 | West | No |
| XOR174 | Canada | NhHe-11:DNA7 | Ruiz-Puerta et al. 2023 | West | No |

Table S4. Summary of genetic sex of historical Atlantic walrus specimens. Specimens are distinguished according to specimen type (tusk ivory or bone), population (based on nuclear DNA assignment, if possible) and find location (Sigtuna or any other location, either trade or control). Genetic sex is determined based on the ratio of X and Y chromosomal coverage (see Table S2) following Dierickx et al. 2025.

| **Speciment type** | | **Tusk ivory** | | | | **Bone (Rostrum or other)** | | | | | |
|---|---|---|---|---|---|---|---|---|---|---|---|
| Population | | Western | | Eastern | | Western | | Eastern | | Unknown | |
| Genetic sex | | ♂ | ♀ | ♂ | ♀ | ♂ | ♀ | ♂ | ♀ | ♂ | ♀ |
| Location | *Sigtuna* | 4 | 3 | 7 | 0 | 1 | 2 | 0 | 0 | 0 | 0 |
| | *Others* | 0 | 0 | 1 | 0 | 31 | 14 | 13 | 6 | 3 | 1 |
